# Mutational Constraint Analysis Workflow for Overlapping Short Open Reading Frames and Genomic Neighbours

**DOI:** 10.1101/2024.07.07.602395

**Authors:** Martin Danner, Matthias Begemann, Florian Kraft, Miriam Elbracht, Ingo Kurth, Jeremias Krause

## Abstract

Understanding the dark genome is a priority task following the complete sequencing of the human genome. Short open reading frames (sORFs) are a group of largely unexplored elements of the dark genome with the potential for being translated into microproteins. The definitive number of coding and regulatory sORFs is not known, however they could account for up to 1-2% of the human genome. This corresponds to an order of magnitude in the range of canonical coding genes. For a few sORFs a clinical relevance has already been demonstrated, but for the majority of potential sORFs the biological function remains unclear. A major limitation in predicting their disease relevance using large-scale genomic data is the fact that no population-level constraint metrics for genetic variants in sORFs are yet available. To overcome this, we used the recently released gnomAD 4.0 dataset and analysed the constraint of a consensus set of sORFs and their genomic neighbours. We demonstrate that sORFs are mostly embedded into a moderately constraint genomic context, but within the gencode dataset we identified a subset of highly constrained sORFs comparable to highly constrained canonical genes.

## Introduction

Variants in the human genome can lead to a variety of pathologies and genome analysis is increasingly used as a basis for clinical decision making(1). While sequencing technologies have improved rapidly in the past years, an extensive analysis of whole genome data lacks behind. The interpretation of genomic data is often limited to the protein coding parts of the genome, which makes up only 1-2 % of the human genome(2). For the far larger part of the genome, sometimes referred to as “Dark Genome”(3, 4), no adequate analysis strategies are available. Part of this “Dark Genome” are genomic elements called “short open reading frames” (sORFs). These sORFs are non-canonical reading frames shorter than the conventionally defined 100 codons, that may overlap with canonical coding regions and might encode for functional microproteins or fulfill regulatory functions(5). sORFs are dispersed across the whole genome and several attempts have been made to classify and to group them into different subcategories(5–11). Our analysis is based on a consensus paper published in 2023 by Mudge et al.(5) in which the used sORF consensus set has been presented and categorized by their genomic localization in comparison to canonical coding as well as non-coding elements. Six categories have been proposed and annotated with a terminology which we have adapted. These six categories are sORFs falling into canonical genes (intORFs), sorfs falling onto long noncoding RNAs (lncRNA-ORFs), sORFs falling into the 5’ (uORF) or 3’ (dORF) untranslated regions (UTR) of canonical genes and sORFs that start in the 5’ (uoORF) or 3’ (doORF) UTR of a canonical gene but reach into the canonical coding region of the gene(5). For some of these sORFs a potential clinical relevance has been demonstrated(12, 13), although the biological role of the majority of sORFs remains elusive. The actual number of predicted sORFs varies within the literature. Conservative estimates used in the gencode sORF consensus set or in another small consensus set provided by Chen et al.(9) contain a few thousand sORFs. This is in contrast to extremely large sORF datasets presented by Neville et al.(10) and Li et al.(11) which contain hundreds of thousands to millions of sORFs. Ultimately, experimental progress will result in the reduction of false positive annotations, although other strategies have been proposed. Of particular importance for our study is the strategy by Jain et al.(16). They suggested using constraint metrics to reduce the number of predicted sORFs, by excluding sORFs which show near to no constraint. While constraint metrics of sORFs that do not overlap with canonical regions have been calculated before, to our knowledge, a constraint analysis for overlapping sORFs, such as those in the gencode dataset, has not been carried out before and provided to the public, particularly using the gnomAD 4.0 dataset. We selected the gencode dataset for our analysis due to several reasons: it contains sORFs which have been found in multiple ribo-seq studies, has redundant sORFs merged, includes sORFs with the canonical start codon ATG, covers both overlapping and non-overlapping sORFs and has undergone review by an international consensus working group.

## Results

### A. Sample size, statistical power and coverage

While sORFs have largely been investigated for evolutionary conservation(5, 11), an extensive quantification for their selective pressure in a large-scale dataset has not been investigated. Reasons for that can be seen in their short length as sORF analysis can be hindered by low statistical power(14). This is particularly noticeable when examining loss-of-function variants which are less likely to occur. The analysis of loss-of-function intolerance in short genomic elements can be complicated when sample size is limited. With the recent gnomAD(15) 4.0 release, these shortcomings can be overcome for some variant types. GnomAD 4.0 contains whole-genome data of 76,215 and whole-exome data from 730,947 individuals which brings up the total number of reference samples to 807,162 individuals. At first, we tested whether all gencode sORFs are contained in the whole-exome samples. This revealed that 4,274 out of the 7,264 gencode sORFs are included in regions present in gnomAD 4.0 exomes. However, not all of them have sufficient coverage for downstream analysis. Therefore, we limited most of our downstream analysis to data contained in the gnomAD genomes.

### B. Analysis of mutational background

The descriptive analysis was conducted on the MANE select subset of coding variants present in genes and sORFs: In their current release the gnomAD genomes contain 4,320,631 unique missense variants, 2,188,653 synonymous variants and 376,271 high impact SNVs (start loss, stop loss, stop gain and frameshift variants) located in canonical genes. Considering the same MANE select transcripts, the gencode sORFs contain 101,445 missense variants, 44,118 synonymous variants and 19,441 high impact variants. The distribution is visualized in Figure 1. Canonical genes and sORFs show a similar distribution of variants with the majority of variants being missense variants, followed by synonymous variants. Protein truncating variants like frameshift and stop-gain variants are a minority, although the gencode sORFs show a noticeably higher amount of high impact variants like frameshift variants and start loss variants.

**Fig. 1.**
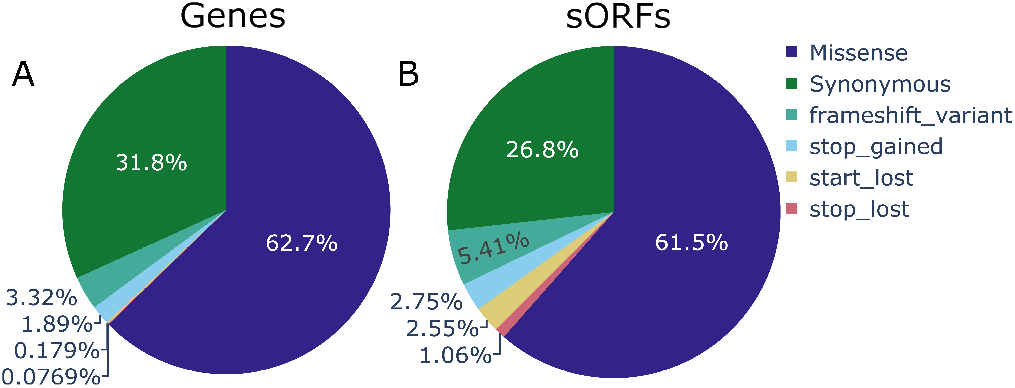
The distribution of variants present in the gnomAD 4.1.0 genomes A) Variant distribution for all MANE select transcripts (genes); B) Variant distribution for all Mane select transcripts (sORFs)

### C. Estimation of sORF constraint utilizing regional genomic constraint

As a next step we analysed whether sORFs are generally intolerant against single nucleotide variants (SNVs). As a first step we annotated the sORFs and for comparison three groups of non-coding genomic elements (lncRNAs, miRNAs and snoRNAs) with the recently published Gnocchi score(17). The distribution of these four elements can be seen in figure 2. As demonstrated previously by Chen et al.(17)a noticeable portion of miRNAs located on autosomes passes the cut off value of 4 (15.4%), while only the minority of lncRNAs show a highly constrained Gnocchi score (2%). To our knowledge snoRNAs were not previously analysed using the Gnocchi score; they show a Gnocchi score comparable with that of lncRNAs. Only 1.5% of snoRNAs fall into the highly constrained percentile of the genome. In comparison, more sORFs show a higher constraint according to the Gnocchi score which places 7% into the most constrained percentile. Subsequently we calculated a constraint metric with a higher resolution, considering only the coding bases of the predicted sORFs. Using the genome data from gnomAD 3.0 / gnomAD 4.0, we calculated the SNV observed/expected upper bound fraction for SNVs falling into sORFs. We termed this value the SNVOEUF, and it is an adaption of the loss-of-function observed/expected upper bound fraction (LOEUF) score proposed by Karczewski et al.(14) and the missense observed/expected upper bound fraction (MOEUF) score put forward by Jain et al.(16).

**Fig. 2.**
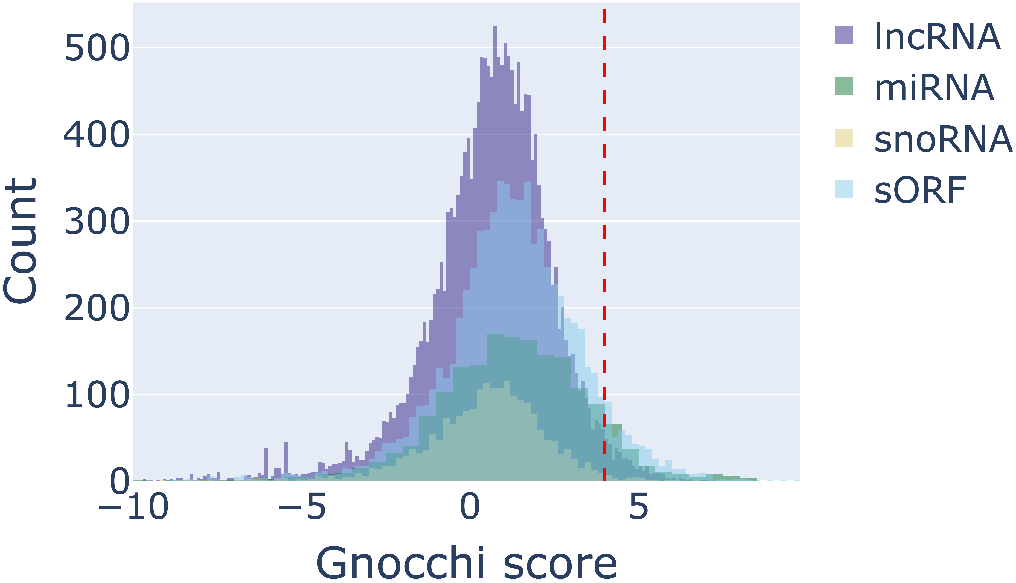
Distribution of paired gnocchi scores of lncRNAs, snoRNAs, miRNAs and sORFs. The cut off vlaue of 4 (here depicted by a vertical red line) marks the border of the most constrained percentile of the genome. The Gnocchi scores were tanken from supplementary Dataset 3 from the Gnocchi score paper(17).

Considering the gnomAD genome data, only 15 sORFs from the gencode dataset have less than 10 expected SNVs and were therefore discarded in the SNVOEUF analysis. The distribution of the SNVOEUF values can be seen in figure 3. Like the Gnocchi score, the SNVOUEF does not allow any assumptions about the coding potential of the analysed region. Instead, it provides an estimate of the overall constraint of the region for SNVs. For downstream analysis we compared the SNVOEUF score to the SNVOEUF score of MANE select transcript filtered coding genes and the SNVOEUF score of the UTR regions corresponding to these MANE select transcript filtered gene list. As visible in figure 3 only a small number of sORFs (53/7,249) passes the cut off value for highly constrained regions, if the highest constraint decile of the SNVOEUF of the UTRs is taking as cut off reference. When compared to genes, 700/7,249 sORFs fall into the highest constraint decile of the SNVOEUF of coding gene regions. The Gnocchi score and SNVOEUF do not correlate well (Kendall Rank Correlation Coefficient=- 0.07, p=6.44e-20). This weak correlation was confirmed in a repeated experiment, in which only the SNVOEUF value of sORFs that fall exactly into a Gnocchi interval was considered (Kendall Rank Correlation Coefficient=-0.1, p=5.76e-22). The SNVOEUF distribution plots of the individual sORF subsets can be found in the appendix.

**Fig. 3.**
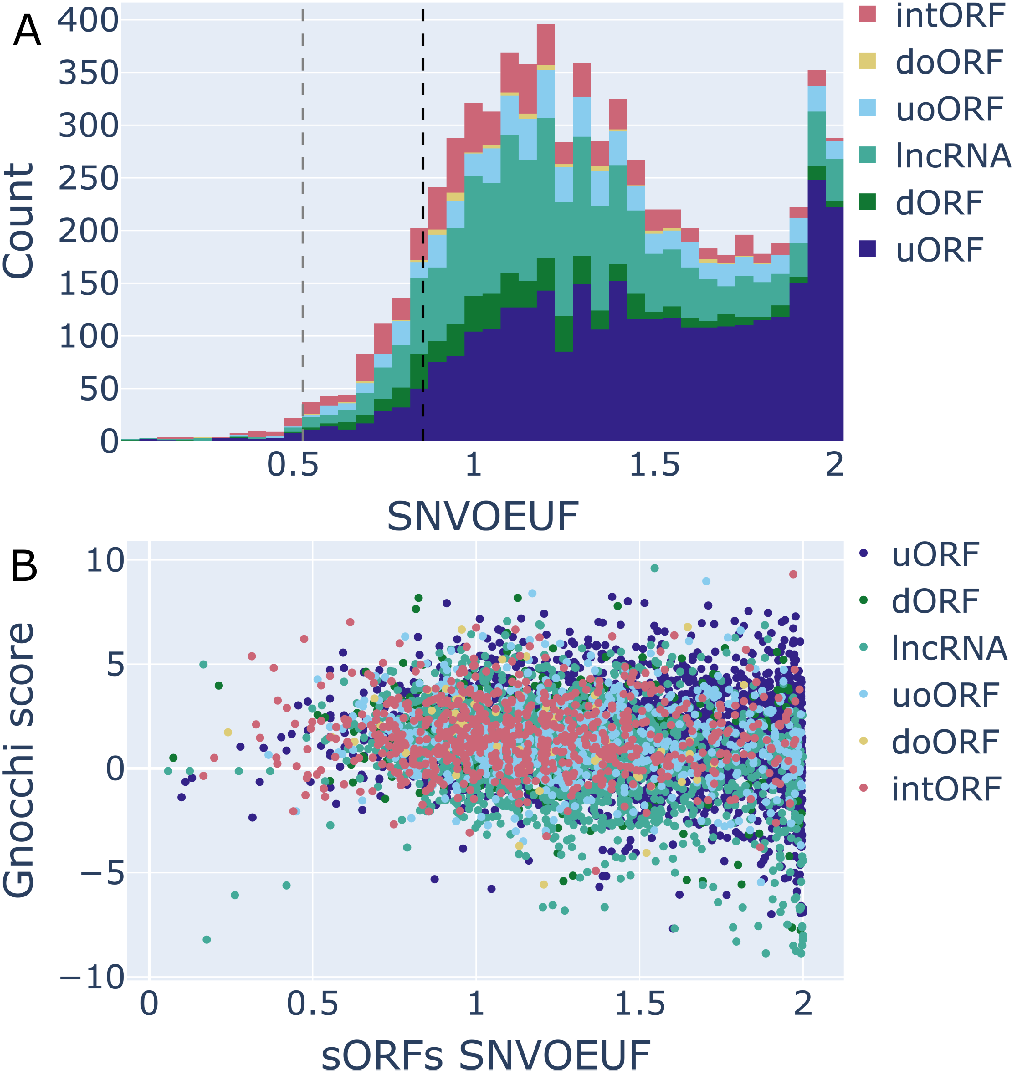
Histogram of the SNV observed/expected upper bound fraction for the different classes of gencode sORFs. We used 0.86 and 0.53 as cut-off values for intolerance. OEUF scores are best interpreted in a continuous manner, but for downstream analysis decile-based filtering is suggested by gnomAD. 0.86 (black vertical line) is the SNVOEUF value of the mane select transcripts of canonical genes. 0.53 (grey vertical line) is the SNVOEUF value of UTRs from mane select transcripts. B) Scatterplot depicting the relation between SNVOEUF values, and the Gnocchi score obtained from the supplementary dataset 3 from the Gnocchi score publication(17).

### D. Most sORFs show a moderate constraint against missense variants

Because sORFs might encode for functional microproteins we additionally calculated a missense constraint score (MOEUF) and loss-of-function constraint score (LOEUF) to unravel the constraint for specific mutational categories, which might be missed when only the SNVOEUF is analysed. Again, we computed the number of expected variants for both classes, to estimate whether reasonable assumptions can be made about a mutational constraint considering the sample size present in gnomAD. Utilizing the genomes from gnomAD 3.0 / 4.0 5,573 out of 7,264 gencode sORFs had equal to or more than 10 expected missense variants. For the same dataset only 2 sORFs had equal to or more than 10 expected loss-of-functions variants, highlighting that despite its size gnomAD is not large enough to make estimates about loss-of-function intolerance using OEUF values for sORFs. Based on the results of the gnomAD flagship paper, which was published using data from 141,456 samples(14), the authors concluded that approximately 75% of genes had a power sufficient for constraint analyses and extrapolated their expected values to estimate a required sample size sufficient for constraint analyses. Since the average sORF is by orders of magnitude shorter than the average canonical gene, we revisited their extrapolation and estimate the required number of sample sizes for LOEUF analyses to be around 5,000,000 samples. While LOEUF analyses therefore are currently out of scope of this paper we restricted our visualization to the distribution of MOEUF scores which can be seen in figure 4. Within the gnomAD genome dataset the sORF MOEUF distribution of (non-overlapping sORFs) differs significantly from the MOEUF distribution of MANE select transcripts of canonical genes when calculated from the genome dataset (Kolmogorov-Smirnov p *<* 0.001). Additionally, a significant difference between these filtered sORF MOEUF values (M_*Rank*_ = 16897.79) and the MOEUF value of the MANE select transcripts of canonical genes (M_*Rank*_ = 10857.72) was observed (Mann-Whitney-U-Test: U=77741709.5, p *<* 0.001). This was further explored by calculating the Vargha and Delaney A measure(33) which returned an estimate of 0.75. The comparison of the two complete distributions can be seen in figure 4B. As visible in figure 4, a small number of sORFs (111 / 5,573) fall into the highly constrained area, which is below the depicted cut off values. To leverage the way larger sample size of the gnomAD 4.0 exome dataset and to estimate the effect of a sample size limitation, we also calculated the MOEUF scores of the previously mentioned 4,274 well covered sORFs included in the gnomAD exome regions. The cutoff value of the highest constrained decile for the MANE select transcripts of the canonical genes was used to study how many sORFs fall into this highly constrained decile. Of the 4,274 sORFs 3,578 had more than 10 expected missense variants and therefore have sufficient data for constraint analysis. Again, within the gnomAD exome dataset the sORF MOEUF distribution (of non-overlapping sORFs) differs significantly from the MOEUF distribution of MANE select transcripts of canonical genes when calculated from the exome dataset (Kolmogorov-Smirnov p *<* 0.001). Similar to the comparison in the genome dataset, a significant difference between these filtered sORF MOEUF values (M_*Rank*_ = 14604.63) and the MOEUF value of the MANE select transcripts of canonical genes (M_*Rank*_ = 9987.03) was observed (Mann-Whitney-U-Tests: U=31096690.5, p *<* 0.001). Again, the Vargha and Delaney A measure(33) was computed and returned an estimate of 0.72. 175 of these 3,578 sORFs fell into the highly constrained decile. Overall, our results are in line with the results presented by Jain et al.(16) who demonstrated that the distribution of MOEUF scores of sORFs tend to be similar to the MOEUF score of less constrained genes. Plots of the MOEUF distributions of individual sORF categories can be found in the appendix.

**Fig. 4.**
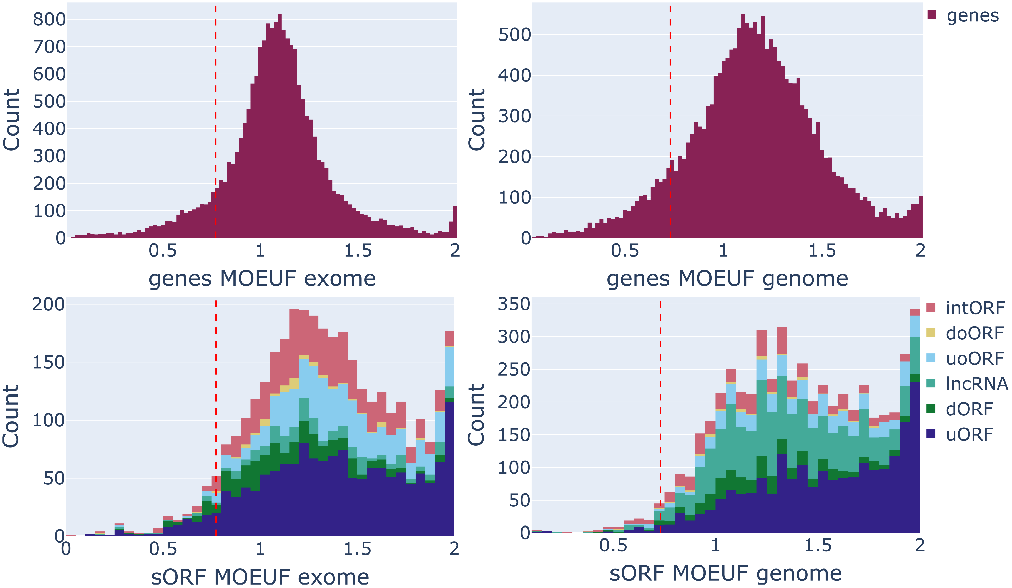
A) MOEUF score of the mane-select transcripts of canonical genes, computed using the data of 730,947 exomes. The red vertical line (0.76) marks the most constrained decile B) MOEUF scores calculated for the MANE select transcripts of the canonical genes, computed using the data of 76,215 genomes. The red vertical line (0.76) marks the most constrained decile C) MOEUF score for the sORFs (which were sufficiently covered in exomes), computed using the data of 730,947 exomes. D) MOEUF score for the sORFs, computed using the data of 76,215 genomes

### E. Most sORFs are neighbhoured by moderately constrained genes

While Jain et al.(16) compared the general distribution of sORFs to the distribution of Refseq genes, to our knowledge the constraint of sORFs has not been previously compared to the directly neighboured genes or genes into which sORFs are embedded and therefore to the genomic background in which sORFs have evolved relatively recently. Therefore, we used the gnomAD exome data to compare the MOEUF values of sORFs to the MOEUF values of their neighboured genes. Figure 5 highlights two points. Firstly, the gencode sORFs tend to be neighboured with genes that are subject to a moderate constraint. Secondly, with a correlation coefficient value of 0.12 (Kendall Rank Correlation Coefficient=0.12, p=5.68e-21) there seems to be only a weak connection between the MOEUF values of sORFs and their neighboured genes. This is further supported if analysed in a pairwise fashion. sORFs (median=1.49) show a significantly higher MOEUF compared to their neighboured genes (median=1.10) (Wilcoxon signed ranked test: W=801505.5, p *<* 0.001). Plots of the individual subcategories of sORFs can be found in the appendix.

### F. Constraint comparison between sORFs and UTRs

1. To further analyse the genomic context of sORFs we investigated the regional constraint of UTRs retrieved from the UTR 2.0 database(20). For this we computed the SNVOEUF score for all UTRs within the UTR 2.0 database assigned to MANE select transcripts, using the data from the gnomAD genomes. Additionally, we differentiated between UTRs that contain gencode sORFs and those that do not. In a similar vein to the MOEUF comparisons, we calculated deciles and picked the maximum SNVOEUF value of the most constrained decile (0.53) as a cut off value for SNV-intolerant UTR regions. As it can be seen in figure 6, UTRs which contain uORFs are dispersed across the SNVOEUF distribution of UTRs. Their distribution is significantly different to the SNVOEUF distribution of UTRs without uORFs (Kolmogorov-Smirnov p *<* 4.14e-52). A significant difference between the sORF containing UTR SNVOEUF values (M_*Rank*_=22516.53) and the SNVOEUF values of UTRs without sORF (M_*Rank*_= 19926.71) was observed (Mann-Whitney-U-Test: U=46219468.0, p *<* 1.06e-29). The subsequently computed Vargha and Delaney A measure returned an estimate of 0.56. To further analyse the UTR regions,

**Fig. 5.**
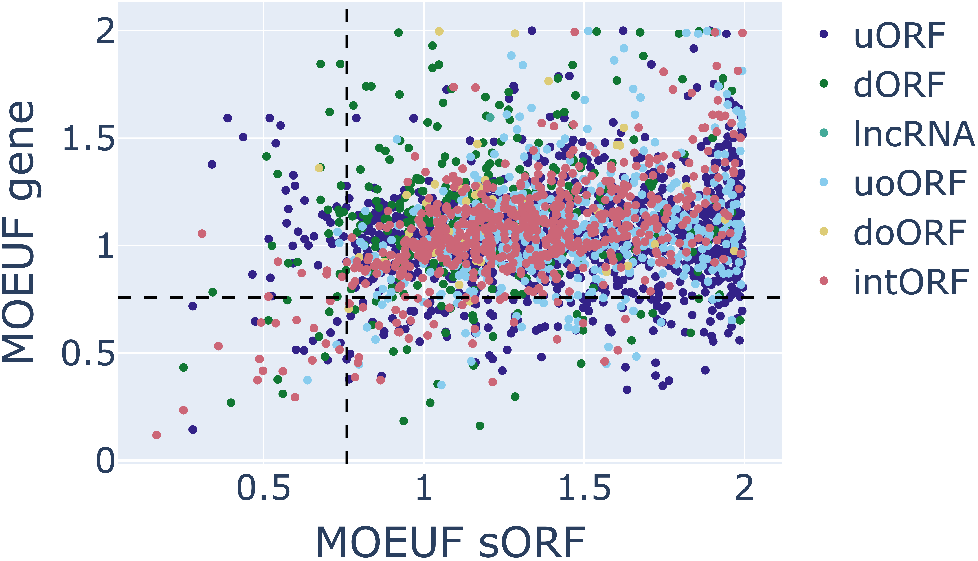
Relationship between the calculated sORF MOEUF and the MOEUF of the neighboring gene, calculated using gnomAD exome data.

**Fig. 6.**
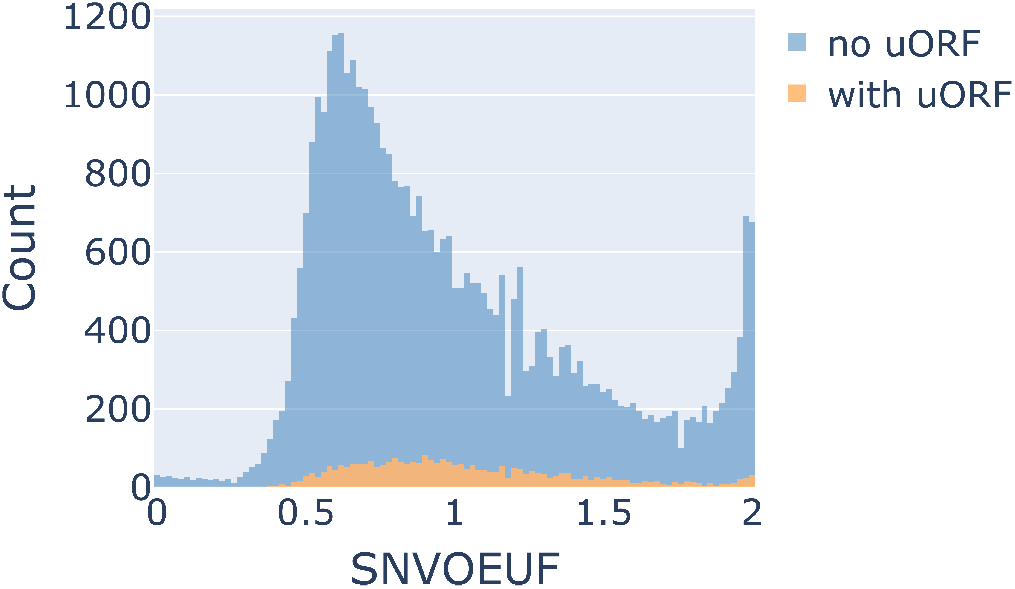
UTROEUF distribution of UTR regions containing uORFs (orange) and without uORFs (blue).

we examined whether canonical genes with multiple UTR regions show regional constraint between their UTRs. As previously introduced, we filtered the annotated UTRs for UTRs with at least 10 expected variants. This reduced the original UTR dataset from 46,216 regions to 40,265 UTRs. Next, we filtered for genes with multiple UTRs, which revealed that out of a total of 18,546 genes, 16,217 genes contain multiple UTRs. To analyse how many of these genes have regional UTR constraint we made use of the previously introduced decile method. We defined a gene as regionally UTR constrained if it contains multiple UTRs which are in separated deciles and delimited by at least two other deciles. Applying this definition 12,610 of the genes contained in the mane select filtered UTR 2.0 database fulfilled this criterion for regional UTR constrain. This is partially explainable by large base length differences observed across the UTRs, as OEUF values are influenced by base sequence length(14). Thus, a correlation analysis showed a strong correlation between the SNVOEUF score and base sequence length (Kendall Rank Correlation Coefficient=-0.46, p *<* 0.001). Being limited by the sample size of gnomAD for small UTRs and to further adjust the analysis for sequence length, we repeated the previous analysis on a sequence length filtered UTR subset, by filtering the 40,265 UTRs for UTRs with a minimum length of 800 bases which corresponds to the mean UTR sequence length of the filtered 40,633 UTRs. Applying these filter criteria, 203 genes have multiple UTR sequences and 107 of these 203 genes have a regional UTR constraint following the definition above. Highlighting the potential relevance of constraint values that incorporate the genomic context of UTRs.

## Methods

### G. Annotation of sORF encoded variants

We utilized the custom annotations feature provided by the ensembl variant effect predictor (VEP)(21, 22) to annotate the gnomAD chromosome reference VCF files for functional consequences in the sORF reading frame. To do so we created a gene transfer format (gtf) file in which we defined the sORF reading frame, by designating the sORF regions from the gencode ribo-seq orf bed file(5) as gene regions and the corresponding block regions as exons. Next to an annotation for functional consequence the general population frequency from gnomAD 4.0 exomes or genomes was added.

### H. Constraint evaluation of sORF encoded variants

For the constraint evaluation of sORF encoding variants, we calculate the observed/expected upper bound fraction (OEUF) for different variant types and name them in correspondence with their variant type (MOEUF for missense variants, LOEUF for loss-of-function variants and SNVOEUF for not further divided SNVs). In brevity, the number of uniquely observed protein truncating variants is compared with the number of expected variants for a given mutation type calculated by a mutational model which assumes a neutral effect of these variants.

For the calculation of observed variants, we followed the recommendations used in the assembly of the gnomAD constraint scores(14). This means that we annotated the gnomAD 4.0 release respecting the sORF context utilizing VEP, then filtered it for the number of unique variants for the mutation type of interest. We only included variants with a general gnomAD frequency less than 0.1 % that passed all filters and had a median depth greater or equal to 1. For all constraint calculations we only considered single nucleotide variants, therefore for loss–of-function variants we only considered start-loss, stop-loss, and stop-gain variants. As a consequence of using VEP(21, 22) for annotation, splice variants are predicted. Considering the limited known information of sORFs and therefore the uncertainty about the occurrence of possible splicing mechanisms in sORFs, we decided to discard variants that were only predicted as splicing variants. This exclusion applied to all analysis, including the loss-of-function analysis.

To calculate the number of expected variants we at first calculated the number of possible variants for which we followed the protocol defined by Karczewski et al.(14). In short, we estimate the number of expected SNVs, missense and synonymous variants by utilizing the mutation rates published in the paper by Chen et al.(17), where the Gnocchi score was proposed. Instead of relying on the Hail-framework previously introduced by Karczewski et al.(14) we reimplemented the workflow by means of Python and Spark. For this, we first extracted the base sequence of the gencode sORFs using biopython(23). Subsequently, we parsed the sequences and calculated the number of possible variants, for the class of interest, by iterating over each coding triplet and calculating the relevant context triplet for each base in the analysed triplet. Adhering to the protocol set by Karczewski et al.(14) we reduced the number of possible variants. Variants were excluded if they originated from bases with a low-quality variant in the gnomAD data, had a high allele frequency (greater or equal to 0.1%) or fell into a region with insufficient coverage (mean depth less than 1). We additionally calculated a sORF mutation rate by summing the mutational rate of each individual base with its corresponding context triplet and the corresponding methylation status. Figure 7 illustrates the workflow schematically. For the final OEUF calculations, we computed the 90% confidence interval (between the 5th and 95th percentiles) for an expected value of a Poisson distribution given an observed count and an expected count.

**Fig. 7.**
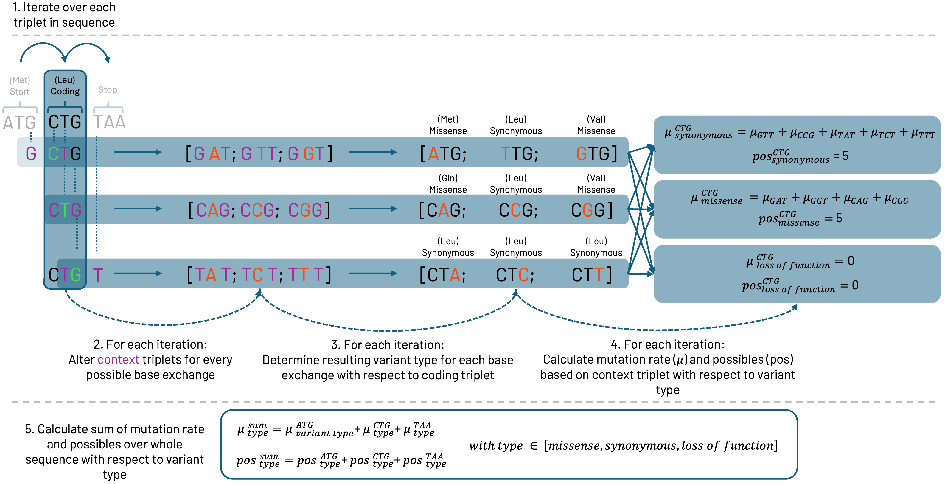
presents a detailed workflow for the calculation of possible variants and mutation rates for a given sequence, based on the type of variant. The iterative process is represented by a sample sequence composed of three codons: the start codon ATG, the triplet CTG, and the stop codon TAA (1). To further elucidate the process, the calculation of the possible variants and mutation rate for the CTG triplet is emphasized. The first step involves identifying the context triplets for each of the three bases (highlighted in green) of the coding triplet. This is done by considering their direct neighboring bases (shown in purple). Afterwards, both the coding and the context triplet are modified to create all possible single nucleotide variants. This is achieved by replacing the bases of the coding triplet (in green) with all other possible bases, resulting in a total of 9 altered triplets (2). Subsequently, the consequence, and thus the variant type, of each altered triplet is determined by evaluating its impact on amino acid translation (3). The mutation rate for a specific variant type of the whole codon is calculated by summing up the mutation rates of the corresponding context triplets (4). Complementary, the count of possible variants for a given variant type is simply the sum of triplets that align with that variant type (4). Finally, the mutation rate and total count of possible variants for the entire sequence are computed by summing up the mutation rates and possible variant counts, respectively, for each variant type across all codons in the sequence (5).

The mathematical formulation of the Poisson probability mass function (pmf) is:

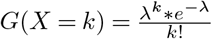

where:

- *λ* is the average rate (expected value)
- k is the actual number of events (observed value)
- e is the base of the natural logarithm
- *G*(*X* = *k*) is the probability of k events occurring in an interval

Here the Poisson pmf is calculated for a range of values from 0 to 2 with steps of 0.001 so that:

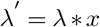

where:

- x is the range value with {*x* : *x* ∈ [0, 2], with *x* = *n* * 0.0001,where *n* ∈ ℕ_0_ and 0 *<*= n *<*= 2000}
- *λ* is the average rate (expected value)

The lower and upper bounds of the expected value are then calculated as the first values in the cumulative distribution where the normalized values are greater than or equal to 0.05 and 0.95 respectively.

The following is a rough mathematical representation of the process:

1. Calculate Poisson pmf for each *λ*^*′*^ for a given expected and observed count
2. Calculate the cumulative sum of these Poisson pmf values
3. Normalise these values
4. Lower Bound = min(*λ*^*′*^ | Normalised Cumulative Poisson pmf *>*= 0.05)
5. Upper Bound = min(*λ* ^*′*^| Normalised Cumulative Poisson pmf *>*= 0.95)

### I. Comparison of OEUF values and defining constraint

OEUF values represent a continuous value which are best analysed as a spectrum, however introducing cut off values for downstream analysis can be helpful. Therefore, following the protocol by Karczewski et al. we binned the resulting OEUF distributions into 10 equally sized bins and took the maximum OEUF value of the most constrained bin as a cut-off value for highly constrained items in this decile. Using this cut off value we further compared sORFs and their OEUF values to other genomic elements.

### J. Matching with Gnocchi Score

In brevity, the Gnocchi score is a constraint metric for which the calculation of the whole genome data from gnomAD 4.0 (76,215 individuals) was analyzed(17).For this the genome was fragmented into 1,000 base spanning intervals. Using a context dependent mutational model for each interval the number of expected variants was calculated and compared to the number of rare SNVs. The resulting ratio was transformed into a z-score and since this score was calculated for every arbitrary region, we paired it with the coordinates of the gencode sORFs located on autosomes(17). Since not all sORFs might encode stable microproteins we wanted to start our analysis with a constraint score in Comparison with three noncoding regions. For this we used the Supplementary Dataset 3 from the recently published Gnocchi score publication (17). This dataset provides genome wide scores at 1 kilobase resolution, calculated iteratively by sliding 100 base pairs. By means of Python the gencode sORFs, the snoDB 2.0(19) snoRNAs, gencode(18) miRNAs and gencode(18) lncRNAs were paired with Gnocchi scores. To be more precise, we located all Gnocchi score Intervals that fell between the start and end coordinates of the genomic element of interest for all autosomes and then the averaged the Gnocchi scores over these intervals. If no proper interval was found containing the genomic element, the two closest Gnocchi score intervals were used to calculate an average Gnocchi score. To have a more direct comparison between the Gnocchi score and the SNVOEUF we subsequently repeated our analysis, by using the non-overlapping Gnocchi dataset (Supplementary Dataset 2 from the Gnocchi score publication(17)) to match the Gnocchi score with sORFs which fall into exactly one Gnocchi score interval. Not mappable sORFs where discarded in this repeated analysis.

### K. Constraint analysis of UTR variants

UTR regions for each MANE select transcript were selected from the UTR 2.0 database(20). We calculated the number of variants in gnomAD 4.0 within said UTR regions and compared them to the number of variants within sORFs located in said UTR region. This was carried out to estimate whether UTR regions containing the sORFs are subjected to a higher degree of selection. The underlying idea is that an UTR containing regulatory elements for the canonical CDS in addition to sORFs which might act as further regulatory elements or encode functional distinct micropeptides, should be subject to a stronger selection than an UTR only fulfilling regulatory purposes. To estimate this effect, we calculated the single nucleotide variant observed/expected upper bound of these UTRs. To analyse a possible correlation between these two values we calculated the Kendall Rank Correlation Coefficient.

### L. Constraint analysis of canonical genes and comparison to sORFs

For the analysis of canonical genes, we filtered the comprehensive VCFs for the current gnomAD 4.0 release for MANE select transcripts using Spark for observed variants with the above-mentioned criteria. We calculated the constraint values in the same fashion as described above and paired the sORFs with their neighboured genes. To analyse a possible correlation between these two values we calculated the Kendall Rank Correlation Coefficient.

### M. Filtering gnomAD exomes for covered sORFs

Utilizing the allele sites published by gnomAD for the exomic regions and the depth summary, we removed possible variants which were insufficiently covered. For this we filtered out sORFs which were not completely contained in the exomic regions file. Additionally, following the gnomAD flagship paper we discarded regions in our analysis that showed a mean coverage of less than 1.

### N. Creation of a whole genome methylation map

In a comparable approach to the gnomAD flagship paper, we obtained the bisulfite whole-genome data provided by the NIH Roadmap Epigenomics Consortium(24). For each genomic position we averaged the methylation fraction across the provided 37 epigenomes from different tissue types and developmental time periods. Afterwards we performed a liftover to hg38 coordinates, using the UCSC chain files and a Python package liftover. Since we used the mutation rates published with the recent Gnocchi constraint score paper, we followingly binned the averaged methylation fractions into 16 bins between 0 and 1. This resulted in the following methylation fraction bins: [0-0.0625, 0.0625-0.125, 0.125-0.1875, 0.1875-0.25, 0.25-0.3125, 0.3125-0.375, 0.375-0.4375, 0.4375-0.5, 0.5-0.5625, 0.5625-0.625, 0.625-0.6875, 0.6875-0.75, 0.75-0.8125, 0.8125-0.875, 0.875-0.9375, 0.9375-1]. We used the corresponding bin of each genomic position to decide which mutation rate to select from the precalculated mutation rates(17) at potentially methylated regions.

### O. Statistics

Most statistical tests were performed using the Python library SciPy(25). For the calculation of the Kendall Rank Correlation Coefficient(26) and the corresponding p-value the SciPy(25) function kendalltau was used. To test the equality of different OEUF distributions a two sample Kolmogorov-Smirnov(27) was performed using the ks_2samp function. To perform the comparison of the mean ranks of different OEUF distribution a Mann-Whitney-U test(28, 29) was performed using Scipys(25) mannwhitneyu function. The mannwhitneyu function from Scipy(25) does not return the calculated mean ranks of the two groups, therefore we utilised the rank function from the Python package pandas to rank the two groups in a concatenated dataframe. Subsequently we summed the ranks of the two groups and calculated an average. For the Mann-Whitney-U test we filtered the sORF dataset for uORFs, dORFs and lncRNA-ORFs to avoid dependencies which would result from using overlapping sORF and coding genomic regions (as present in intORFs, uoORFs and doORFs) for constraint calculation. For the comparison between the sORF MOEUF values and their neighboured gene MOEUF values a Wilcoxon signed ranked test(28, 29) was performed using SciPy(25) and the Wilcoxon function. Multiple hypothesis tests and correlation analysis resulted in p-values not distinguishable from 0 by scipy and R(31). These values were reported as p value *<* 0.001 as suggested by multiple scientific organisations (e.g. the APA). Following the suggestions by Lin, Lucas and Shmueli(30) for statistical inference in large and complex datasets, we accompanied most of our hypothesis tests with visualizations and analysis for effect sizes. We utilised the R package effsize(32) to compute the Vargha and Delaney A measure(33), provided by the VD.A function. The Vargha and Delaney A measure can be described as the probability that a value from group one will be greater than a value from group two(34), thereby allowing a further quantification of the difference between the two groups.

### P. Data Processing with Databricks and Spark

The gnomAD 4.0 VCFs underwent comprehensive processing, including the preparation of positional whitelists, the filtration of designated genomic regions such as MANE transcripts, and observed variant count. This processing was conducted using Databricks and Azure Cloud. The employed Databricks runtime was 14.3 LTS, which encompasses Apache Spark 3.5.0 and Scala 2.12. The computational infrastructure utilized was a multi-node E48d V4 Cluster, equipped with 384 GB memory and 48 cores per node.

## Discussion

Research on sORFs is an emerging field and the discussion of their function in the genome and their biological role is still ongoing. Here we tested different approaches to prioritize and evaluate variants that affect sORFs and neighboured regions, which is fundamental to understanding their role in health and disease.

### Q. sORFs show a similar mutational background to coding regions

sORFs show a similar mutational background to canonical genes, yet they can contain a higher number of high impact variants. This can have multiple explanations. It might be that these regions are not intolerant against loss-of-function variants or that these non-constrained sORFs do not encode functional microproteins. This similarity in distribution, as seen in figure 1, on its own, does not bring sufficient evidence for a potential coding effect or conservation in sORFs, because the distribution might be fully explainable from a probabilistic standpoint, since synonymous and protein truncating variants have less opportunities to occur compared to missense variants. Followingly more complex workflows are required that respect the genomic context and the different effect of variants in differing reading frames.

### R. A streamlined context dependent genomic constraint workflow

We put one of these more complex workflows to the test, by analysing OEUF values for the effect of sequence length, alternating reading frames in overlapping regions and different genomic contexts. We provide evidence for the context sensitivity of these scores, therefore highlighting the importance of carefully mapped genomic constraint maps and the need for clearly defined genomic regions. We further strengthen this point by showing the difference in genomic constraint between sORFs and their closest genomic neighbours and transferring mapped OEUF values onto other regulatory regions, such as UTRs. Thereby we present further evidence for regional constraint differences, even in related regions. This is a concept which is in the process of being adapted in the analysis of coding regions, where it already has been demonstrated that the interpretation of some genes benefits from regional constraint values in comparison to gene wide constraint values(35, 36). For this, we sought to establish a more simplified method in comparison to the gnomAD Hail approach. This streamlined procedure enhances accessibility, particularly for Python-native developers who may not be as well-versed with the Hail framework. It empowers the wider community to calculate constraint metrics for their specific genomic elements of interest. While this approach has been specifically adapted for the use case presented here, it is important to acknowledge that the Hail framework offers a far more extensive range of capabilities. While OEUF values are a useful and versatile tool, their limitations need to be kept in mind, especially when comparing OEUF values from multiple regions to one another. OEUF scores are a one-sided conservative estimation. Low OEUF scores hint towards a higher level of constraint, while high OEUF scores can either depict a region which is only weakly constrained or signal a small sample size with limited explanatory power. The limited explanatory power is a present issue, even today with datasets like gnomAD. Our analysis of the mutational background in gnomAD and the calculation of the different constraint metrics demonstrate that the currently available sample size in gnomAD is insufficient to fully analyse the constraint level of sORFs. Especially the analysis of loss-of-function intolerance requires sample sizes which are by a magnitude larger than the current gnomAD release. As previously stated, small genomic regions suffer the most from the limited sample size since they already have a small number of mutational opportunities. Present metrics like the Gnocchi score try to reduce this problem by binning the genome in relatively large regions of 1,000 bases which are not mapped to the exact genomic architecture. We demonstrate that the OEUF value of SNVs in sORFs do not correlate well with the matched Gnocchi score of the sORFs. This might be explained by several possibilities. The first possibility is that the Gnocchi score bins relatively large regions of interest. sORFs only make up a small portion of these intervals and tend to fall within regions overlapping with canonical regions and regulatory elements. Consequently, sORFs might just be another factor under many more that influences the Gnocchi score. Highlighting the importance of context sensitive genomic constraint maps and highlighting that the Gnocchi score might benefit from a higher resolution, a suggestion also mentioned in the original Gnocchi score paper(17). On the other hand, our SNVOEUF calculation might also suffer from the short sample size. Large sample sizes could reveal a correlation between the Gnocchi score and the SNVOEUF which might be masked by the fact that the sORF regions are just smaller and therefore tend to be more influenced by the small sample size, than the larger gnocchi score intervals. Thereby highlighting the interaction of OEUF values, genomic context and sequence length and the need for expanding the presented workflow to larger datasets than gnomAD 4.0.

### S. A subset of the sORF gencode dataset is highly constrained

While the Gnocchi score and the SNVOEUF score can give a general overview of the constraint, additional metrics are required to analyse the coding potential of sORFs. Our analysis using the gnomAD genomes revealed that a portion of sORFs has a similar constraint to highly constrained genes with a similar OEUF value. While repeating this analysis with gnomAD exome data, we interestingly noticed a shift in the MOEUF distribution towards a smaller MOEUF value. We were surprised to see that the number of highly constrained sORFs increased. This could indicate that the current sample size for the gnomAD genomes might still be insufficient with respect to sORFs. As a result, this could potentially even lead to an underestimate of constraint in current datasets. These highly constrained sORFs, although a minority, might be of special interest in terms of potential clinical relevance since they show a similar constraint to the most constrained coding regions of the canonical genes. To our surprise intORFs were a minority in these highly constrained regions, which might highlight that these highly constrained regions are not purely ranked as highly constrained because of an overlap with canonical coding regions. When compared to the neighbouring genes, which is used in the nomenclature of the gencode sORF set, we were not able to witness a correlation for both, gnomAD genome and exome data. This might be partially explained by the fact that the gnomAD constraint values are currently calculated gene wide. Some of these genes might have regional constraints, which will only be uncovered if a regional constraint calculation (exon wide or domain wide) is performed. To fully understand the connection between sORFs and neighbouring genes it would be beneficial to analyse the sORFs and their closest neighbouring gene element, annotated with regional constraint. This pattern of moderate constraint was continued when we compared uORFs with the constraint scores of UTR regions. This highlighted that uORFs evolved across a broad spectrum of constrained UTRs but were not enriched in highly constrained UTRs. Taken together we hypothesize that this moderately constrained background of genomic regions might have been a necessary condition for sORFs to evolve. This is supported by literature(5) as some sORFs are described as phylogenetically young and this background likely provides a balance between room for some genomic change and constraint.

## Conclusion

We implemented a constraint calculation workflow in Python and Spark, comparable with that previously presented by the team from gnomAD. This enabled us to analyse coding and noncoding genomic regions. We benchmarked this workflow on gnomAD 4 exome and gnomAD 3/4 genome data. Utilizing this workflow, we predicted OEUF values for a high consensus set of sORFs which to our knowledge has not been done previously on this scale and compared it to the genomic background of sORFs. We demonstrated the moderate constraint involving most sORFs in our dataset but highlight that a minority of sORFs is constrained in a similar way to canonical genes, highlighting the need for further research in this emerging field. Our calculations establish a measure for predicting the impact of sORF variants and identify a subset of maximally intolerant sORFs within the human genome. This bioinformatic workflow establishes the basis for the further exploration and prioritization of sORF variants and towards a better understanding of the dark genome.

## Supporting information

sORF Annotation scores

## CODE AVAILABILITY

The predicted constraint values from the genome and exome data for the sORFs, UTRs and mane values are available as supplementary dataset accompanying this article. The gnomAD constraint table and the gnomAD VCFs are publicly available on the gnomAD website https://gnomad.broadinstitute.org/downloads. The mutation rates were extracted from the supplementary datasets of gnomAD gnocchi Score paper(17). The epigenomes are publicly available on the NIH epigenomics website https://www.ncbi.nlm.nih.gov/geo/roadmap/epigenomics/. The gencode sORFs were taken from the corresponding project website https://www.gencodegenes.org/pages/riboseq_orfs/. The snoRNAs were taken from the snoDB website https://bioinfo-scottgroup.med.usherbrooke.ca/snoDB/. The UTR data was obtained from the corresponding github page https://github.com/BioinfoUNIBA/UTRdb.

## CODE AVAILABILITY

The Python code used to calculate the OEUF scores will be made availabe upon publication.

## AUTHOR CONTRIBUTIONS

JK, IK, ME, MB and FK conceived the Idea. JK and MD implemented the constraint workflow in Python and Spark. JK and MD parsed gnomAD, the gencode riboseq ORFs and the UTR database. JK and MD computed the OEUF metrics, the comparison to the canonical OEUF values and created the plots. JK, MD and MB wrote the manuscript. JK and MD wrote tests and performed validations. IK, ME, MB, FK and JK provided supervision. All authors were involved in reviewing of the transcript and gave their consent for the submission of the final version.

## ACKNOWLEDGEMENTS

This research project was funded by the START-Program of the Faculty of Medicine, Uniklinik RWTH-Aachen. I.K. is supported by the Deutsche Forschungsgemeinschaft (DFG) (KU 1587/6-1, KU 1587/9-1, KU 1587/10-1, KU 1587/11-1) I.K. receives funding from the European Union’s Horizon 2020 research and innovation programme under the EJP RD COFUND-EJP N° 825575. We would like to extend our sincere gratitude to Jan Höllmer and Lars Perchalla whose support in the early stages was crucial to the inception of this project and the research outcomes presented in this report. Furthermore, We thank Ricardo Henriques and his group for the distribution of their biorxiv latex template via overleaf https://de.overleaf.com/latex/templates/henriqueslab-biorxiv-template/nyprsybwffws.

## Supplementary Note 1: Unravelling the constraint of the individual sORF classes

Additionally to visualising the combined constraint values across all sORF classes, we visualised and analysed the individual sORF classes on their own and provide the results of this analysis in this supplementary material.

### A. SNVOEUF distributions for the individual sORF classes

We began this Analysis by listing the individual SNVOEUF distributions of the sORFs, as calculated using the gnomAD genome data.

**Fig. 8.**
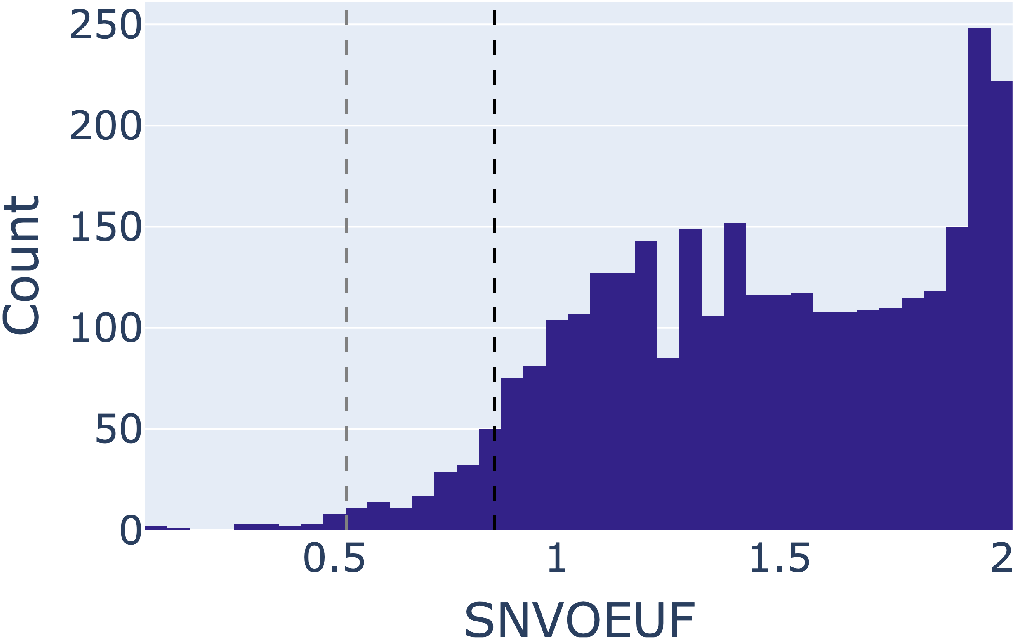
SNVOEUF distribution for the uORF sORF class, calculated using the gnomAD genome data. Decile comparison was performed as introduced previously. 0.86 (black vertical line) is the SNVOEUF value of the mane select transcripts of canonical genes. 0.53 (grey vertical line) is the SNVOEUF value of UTRs from mane select transcripts.

**Fig. 9.**
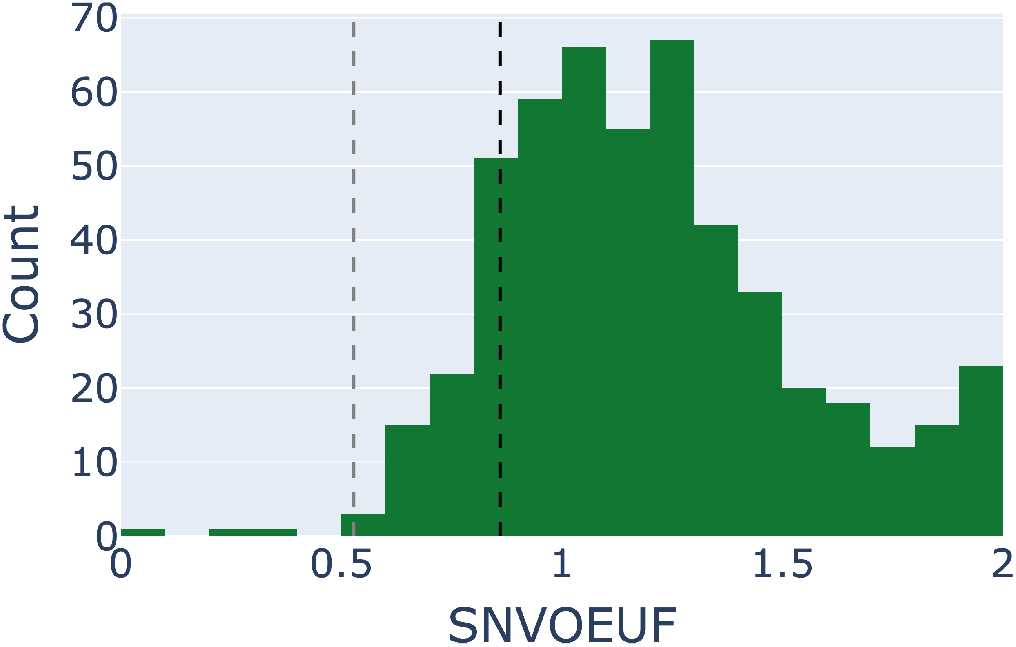
SNVOEUF distribution for the dORF sORF class, calculated using the gnomAD genome data. Decile comparison was performed as introduced previously. 0.86 (black vertical line) is the SNVOEUF value of the mane select transcripts of canonical genes. 0.53 (grey vertical line) is the SNVOEUF value of UTRs from mane select transcripts.

**Fig. 10.**
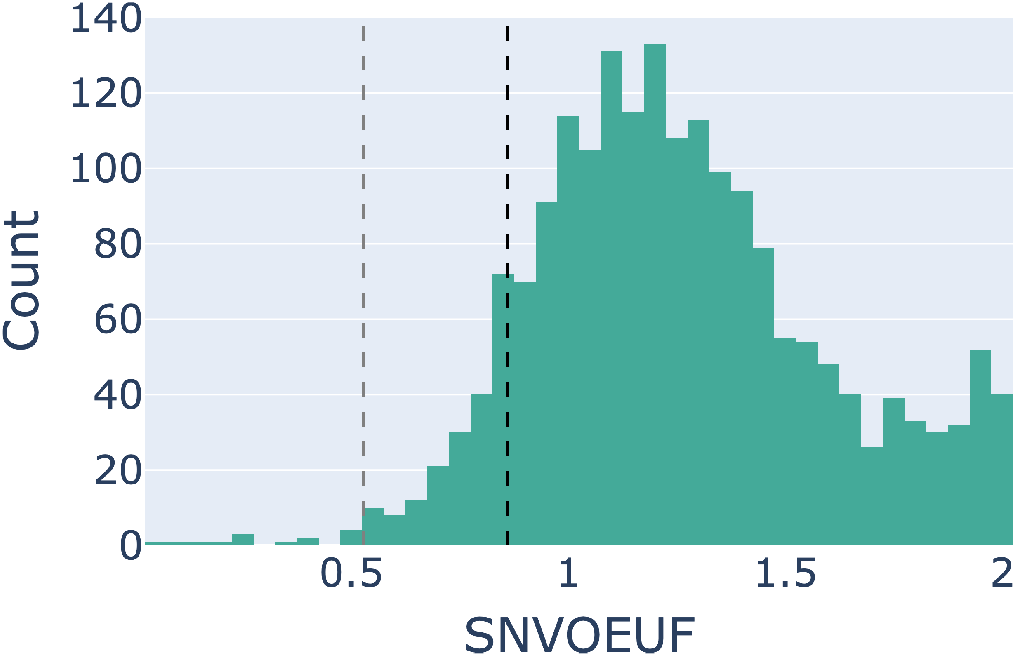
SNVOEUF distribution for the sORF class located on lncRNAs, calculated using the gnomAD genome data. Decile comparison was performed as introduced previously. 0.86 (black vertical line) is the SNVOEUF value of the mane select transcripts of canonical genes. 0.53 (grey vertical line) is the SNVOEUF value of UTRs from mane select transcripts.

**Fig. 11.**
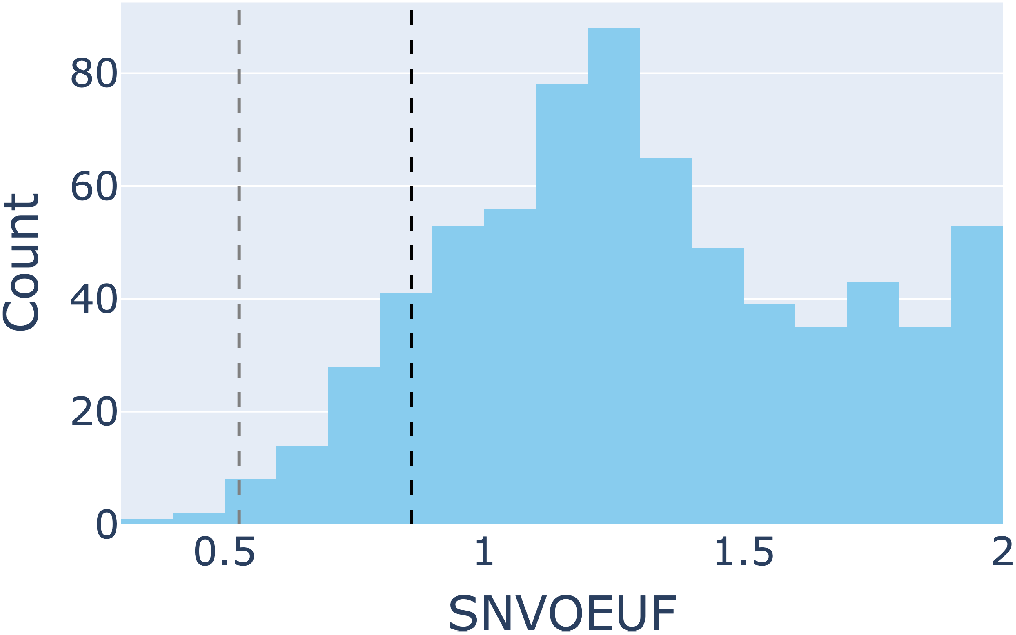
SNVOEUF distribution for the uoORF sORF class, calculated using the gnomAD genome data. Decile comparison was performed as introduced previously. 0.86 (black vertical line) is the SNVOEUF value of the mane select transcripts of canonical genes. 0.53 (grey vertical line) is the SNVOEUF value of UTRs from mane select transcripts.

**Fig. 12.**
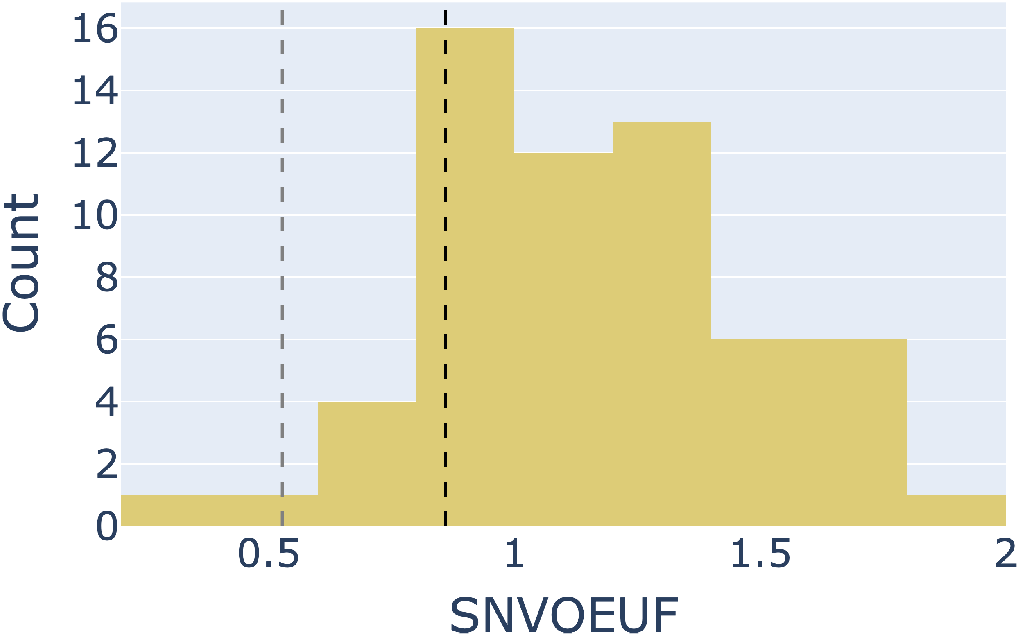
SNVOEUF distribution for the doORF sORF class, calculated using the gnomAD genome data. Decile comparison was performed as introduced previously. 0.86 (black vertical line) is the SNVOEUF value of the mane select transcripts of canonical genes. 0.53 (grey vertical line) is the SNVOEUF value of UTRs from mane select transcripts.

**Fig. 13.**
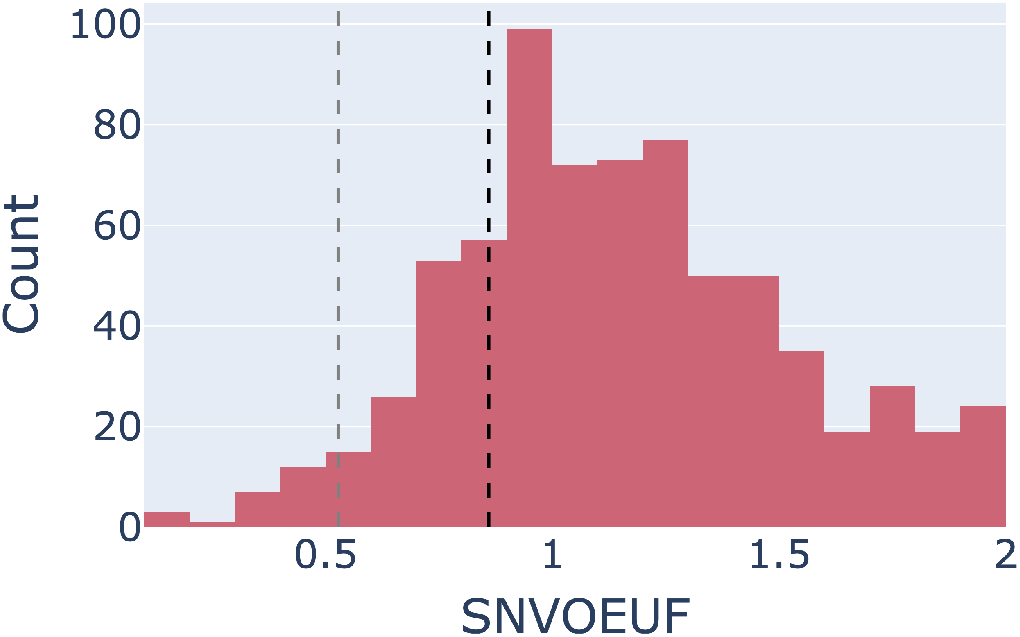
SNVOEUF distribution for the intORF sORF class, calculated using the gnomAD genome data. Decile comparison was performed as introduced previously. 0.86 (black vertical line) is the SNVOEUF value of the mane select transcripts of canonical genes. 0.53 (grey vertical line) is the SNVOEUF value of UTRs from mane select transcripts.

### B. MOEUF distributions for the individual sORF classes (genome data)

Subsequently we listed the MOEUF distributions for the individual sORF classes using the gnomAD genome data.

**Fig. 14.**
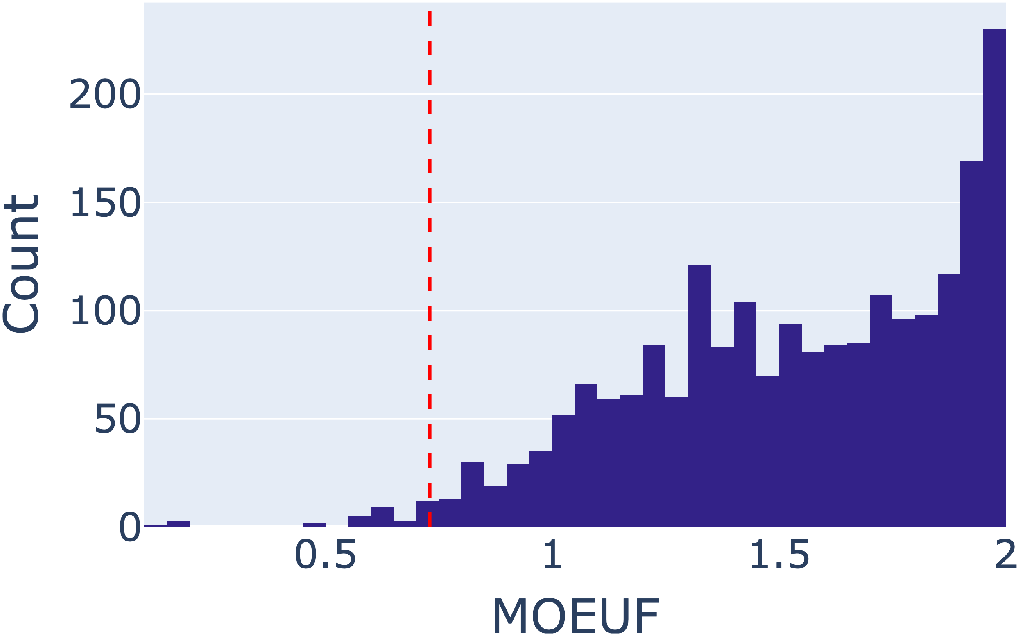
MOEUF distribution for the uORF sORF class, calculated using the gnomAD genome data. Decile comparison was performed as introduced previously. The red vertical line (0.76) marks the most constrained decile of the MOEUF value of OEUF scores calculated for the MANE select transcripts of the canonical genes using the gnomAD genome data.

**Fig. 15.**
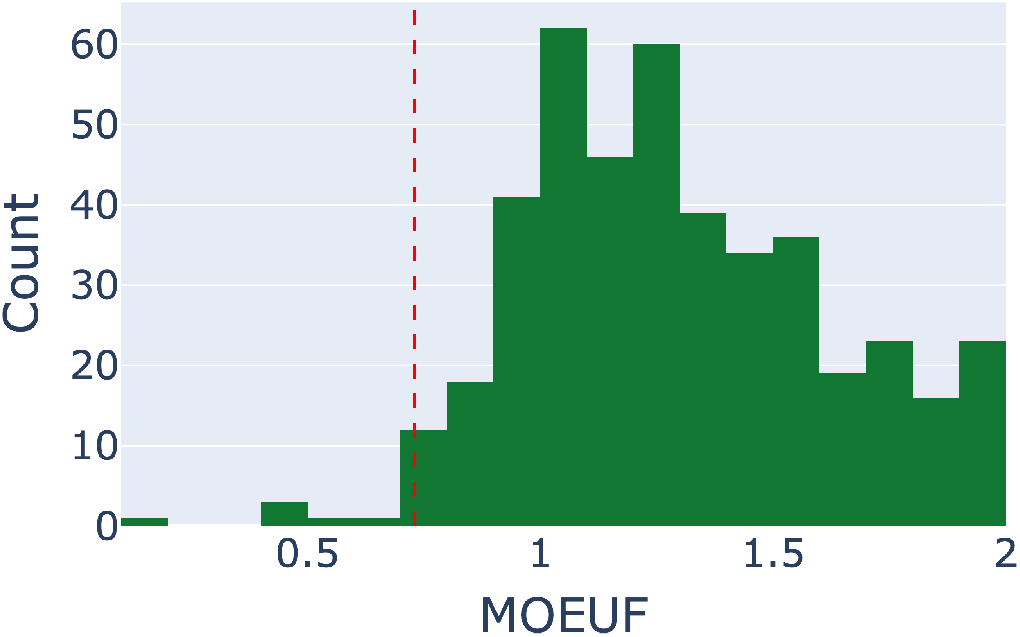
MOEUF distribution for the dORF sORF class, calculated using the gnomAD genome data. Decile comparison was performed as introduced previously. The red vertical line (0.76) marks the most constrained decile of the MOEUF value of OEUF scores calculated for the MANE select transcripts of the canonical genes using the gnomAD genome data.

**Fig. 16.**
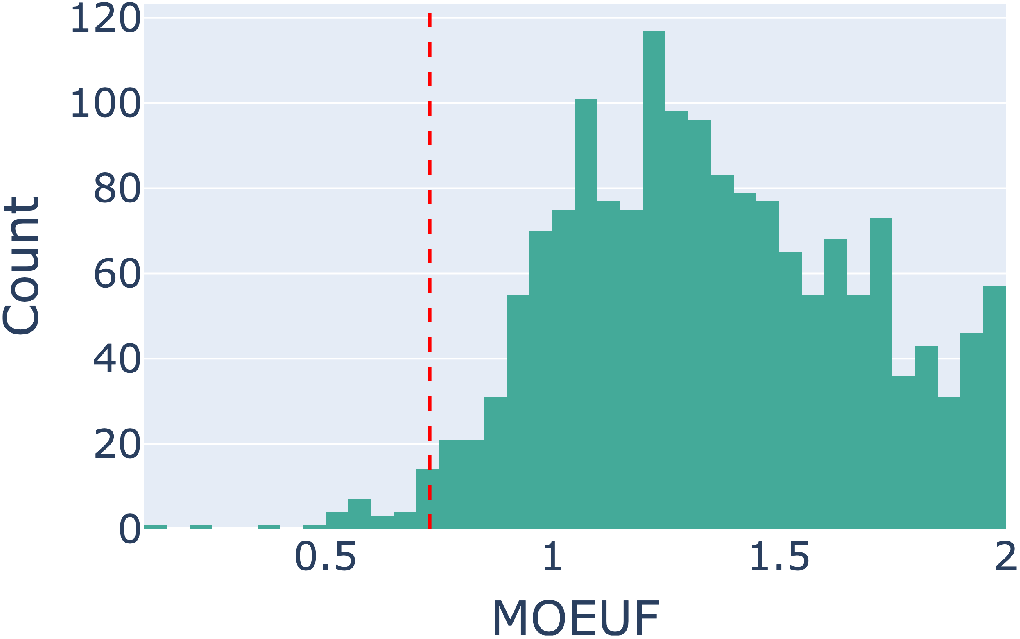
MOEUF distribution for the sORFs located on lncRNAs, calculated using the gnomAD genome data. Decile comparison was performed as introduced previously. The red vertical line (0.76) marks the most constrained decile of the MOEUF value of OEUF scores calculated for the MANE select transcripts of the canonical genes using the gnomAD genome data.

**Fig. 17.**
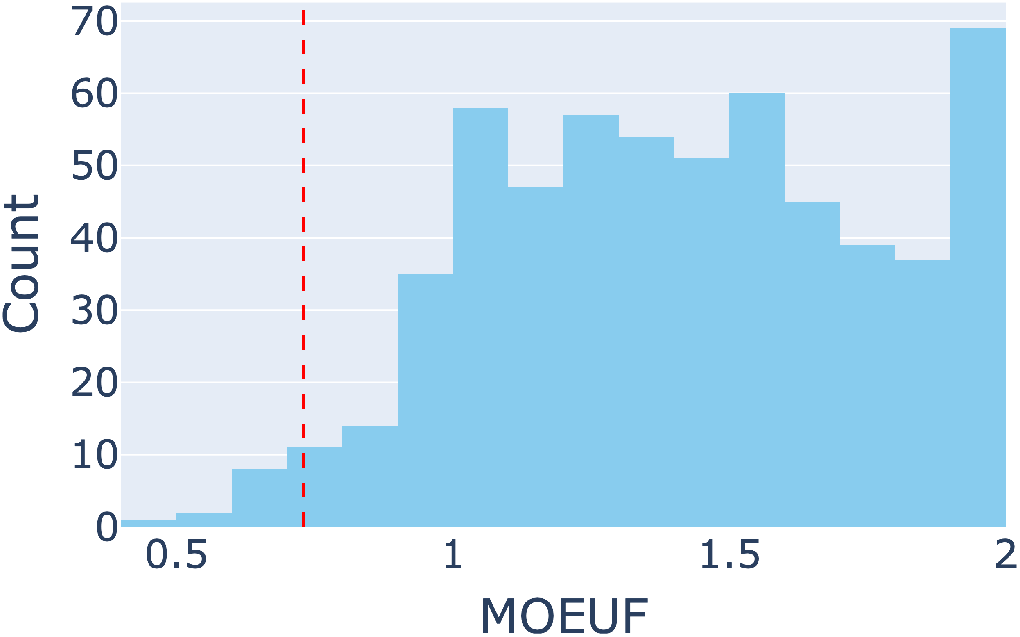
MOEUF distribution for the uoORF sORF class, calculated using the gnomAD genome data. Decile comparison was performed as introduced previously. The red vertical line (0.76) marks the most constrained decile of the MOEUF value of OEUF scores calculated for the MANE select transcripts of the canonical genes using the gnomAD genome data.

**Fig. 18.**
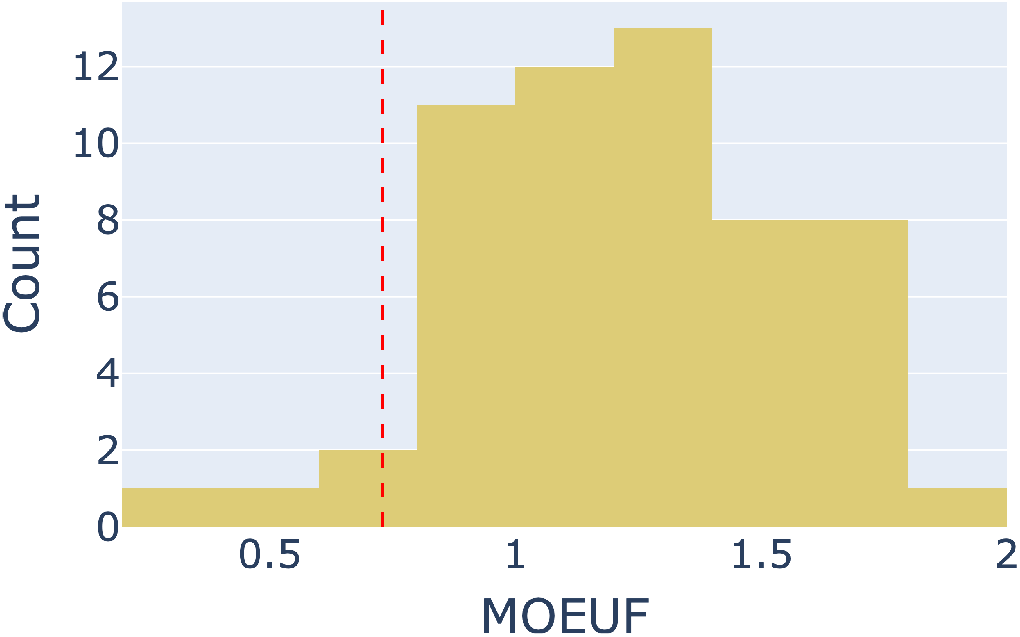
MOEUF distribution for the doORF sORF class, calculated using the gnomAD genome data. Decile comparison was performed as introduced previously. The red vertical line (0.76) marks the most constrained decile of the MOEUF value of OEUF scores calculated for the MANE select transcripts of the canonical genes using the gnomAD genome data.

**Fig. 19.**
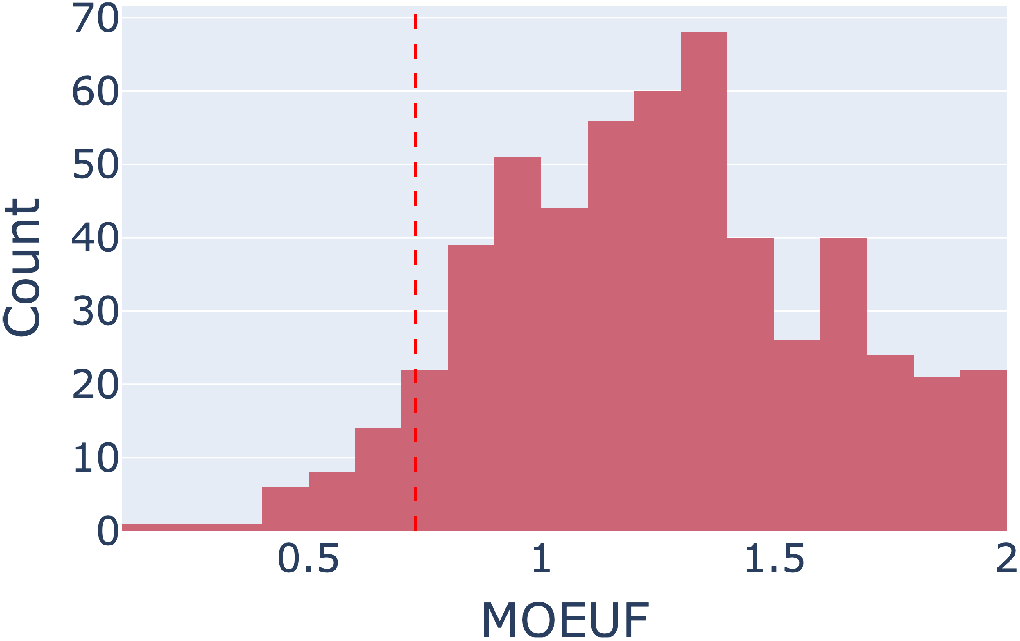
MOEUF distribution for the intORF sORF class, calculated using the gnomAD genome data. Decile comparison was performed as introduced previously. The red vertical line (0.76) marks the most constrained decile of the MOEUF value of OEUF scores calculated for the MANE select transcripts of the canonical genes using the gnomAD genome data.

### C. MOEUF distributions for the individual sORF classes (exome data)

Followingly, we listed the MOEUF distributions for the individual sORF classes using the exome genome.

**Fig. 20.**
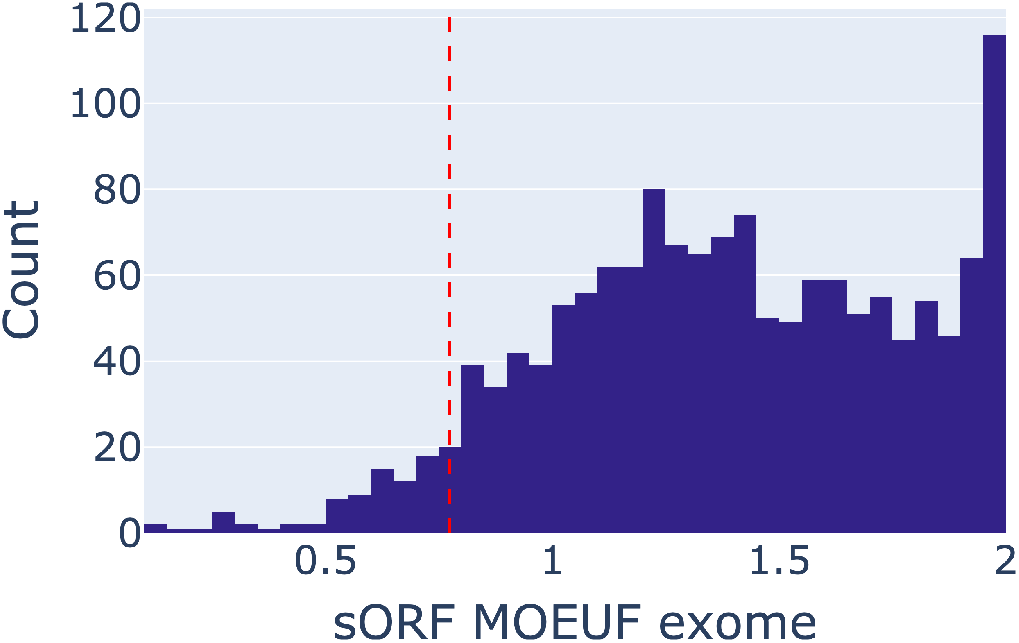
MOEUF distribution for the uORF sORF class, calculated using the gnomAD exome data. Decile comparison was performed as introduced previously. The red vertical line (0.73) marks the most constrained decile of the MOEUF value of OEUF scores calculated for the MANE select transcripts of the canonical genes using the gnomAD exome data.

**Fig. 21.**
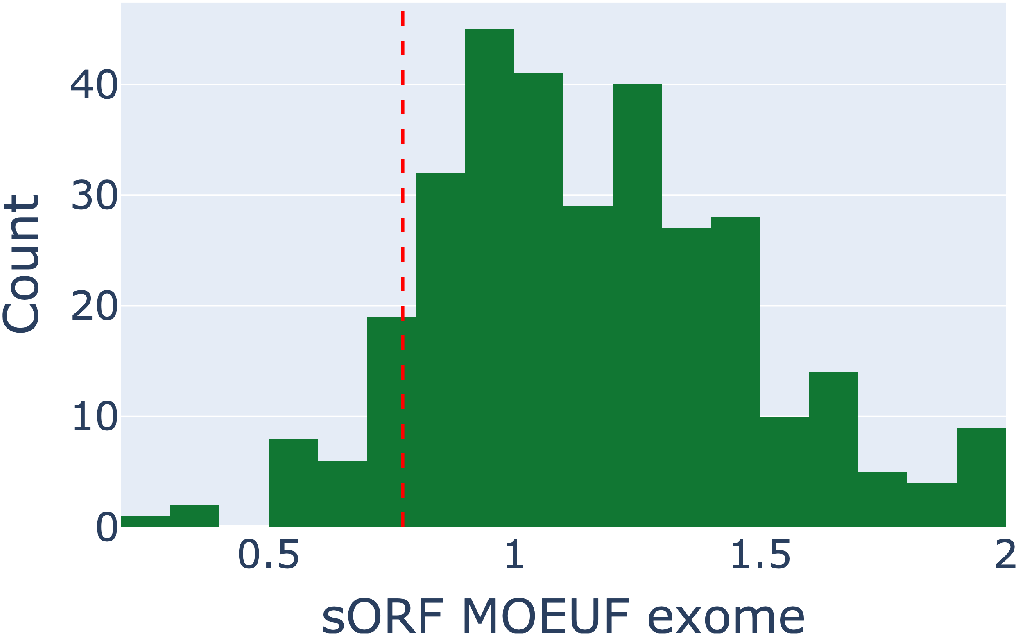
MOEUF distribution for the dORF sORF class, calculated using the gnomAD exome data. Decile comparison was performed as introduced previously. The red vertical line (0.73) marks the most constrained decile of the MOEUF value of OEUF scores calculated for the MANE select transcripts of the canonical genes using the gnomAD exome data.

**Fig. 22.**
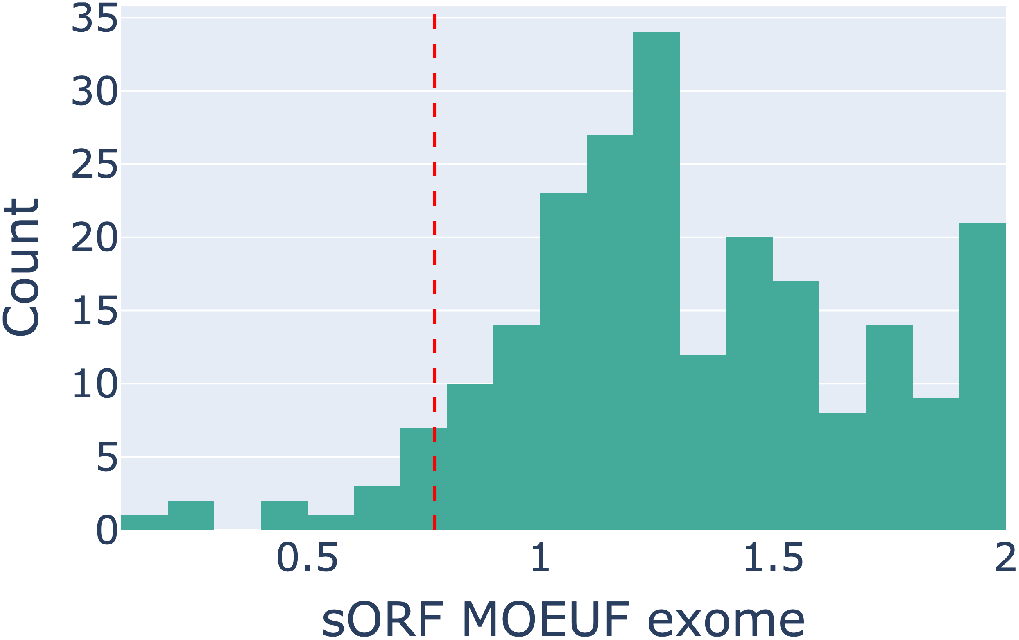
MOEUF distribution for the dORF sORF class, calculated using the gnomAD exome data. Decile comparison was performed as introduced previously. The red vertical line (0.73) marks the most constrained decile of the MOEUF value of OEUF scores calculated for the MANE select transcripts of the canonical genes using the gnomAD exome data.

**Fig. 23.**
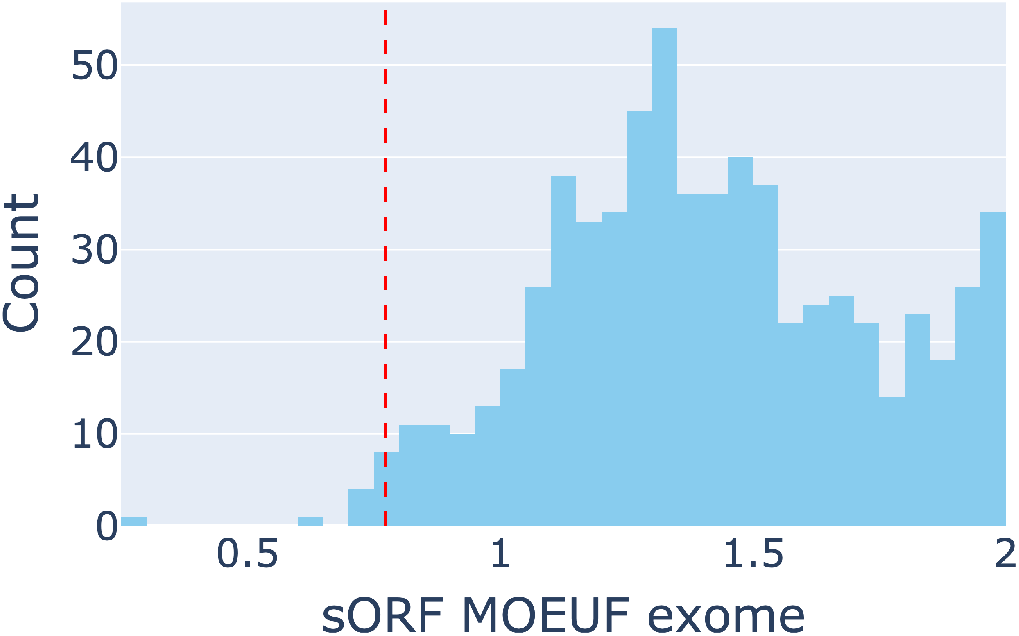
MOEUF distribution for the dORF sORF class, calculated using the gnomAD exome data. Decile comparison was performed as introduced previously. The red vertical line (0.73) marks the most constrained decile of the MOEUF value of OEUF scores calculated for the MANE select transcripts of the canonical genes using the gnomAD exome data.

**Fig. 24.**
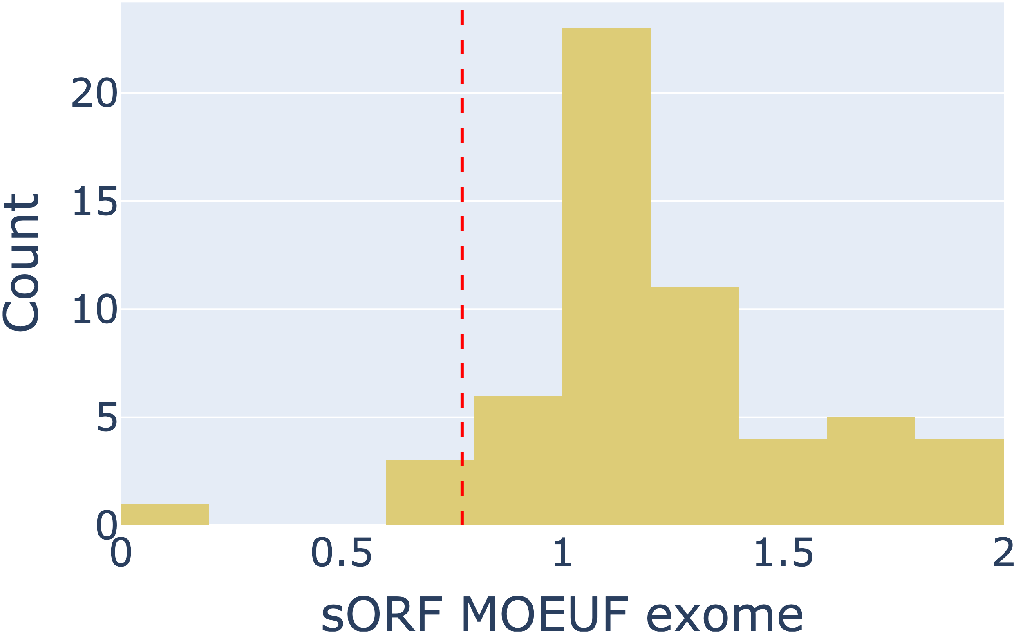
MOEUF distribution for the dORF sORF class, calculated using the gnomAD exome data. Decile comparison was performed as introduced previously. The red vertical line (0.73) marks the most constrained decile of the MOEUF value of OEUF scores calculated for the MANE select transcripts of the canonical genes using the gnomAD exome data.

**Fig. 25.**
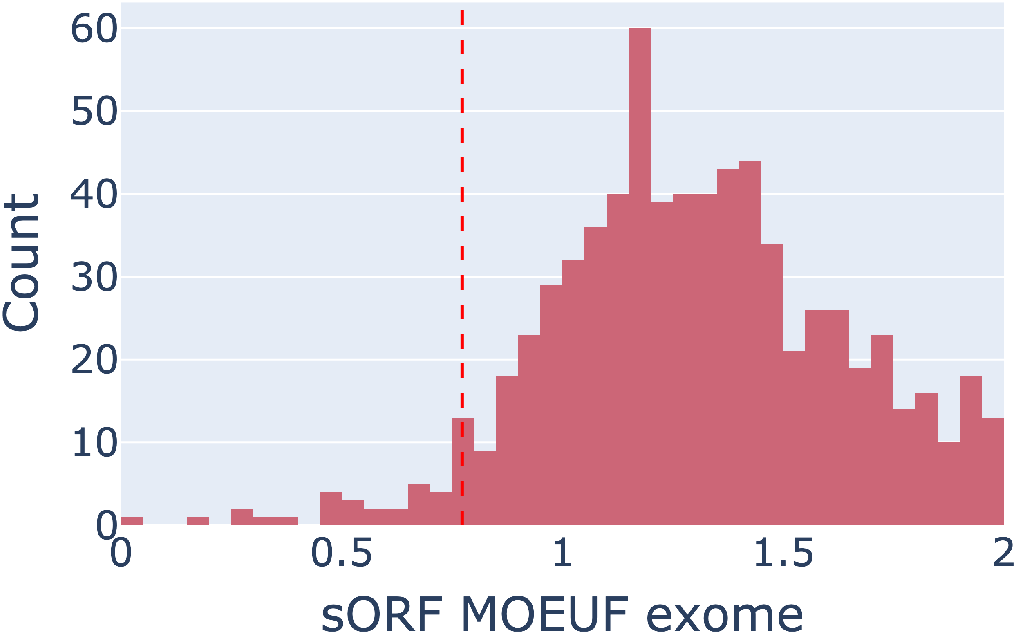
MOEUF distribution for the dORF sORF class, calculated using the gnomAD exome data. Decile comparison was performed as introduced previously. The red vertical line (0.73) marks the most constrained decile of the MOEUF value of OEUF scores calculated for the MANE select transcripts of the canonical genes using the gnomAD exome data.

### D. Individual sORF class comparison with the Gnocci Score

Additionally, to the summary plot found in the article, we provide individual plots for comparison, in which we compare the SNVOEUF score of the individual sORF classes with the Gnocchi Score.

**Fig. 26.**
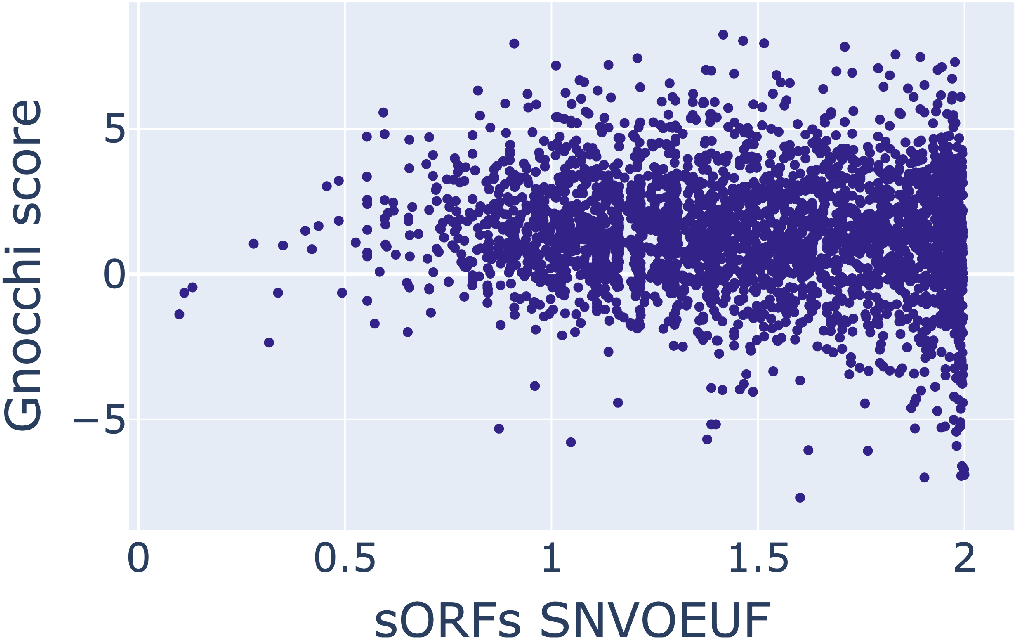
Comparison between SNVOEUF of uORFs calculated using gnomAD genomes and the Gnocchi Score.

**Fig. 27.**
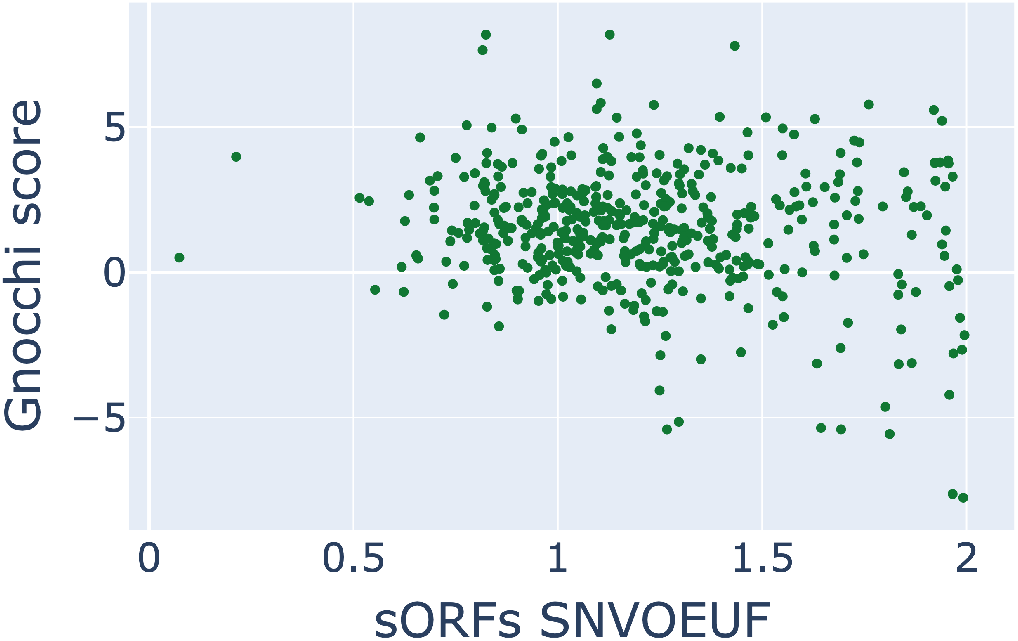
Comparison between SNVOEUF of dORFs calculated using gnomAD genomes and the Gnocchi Score.

**Fig. 28.**
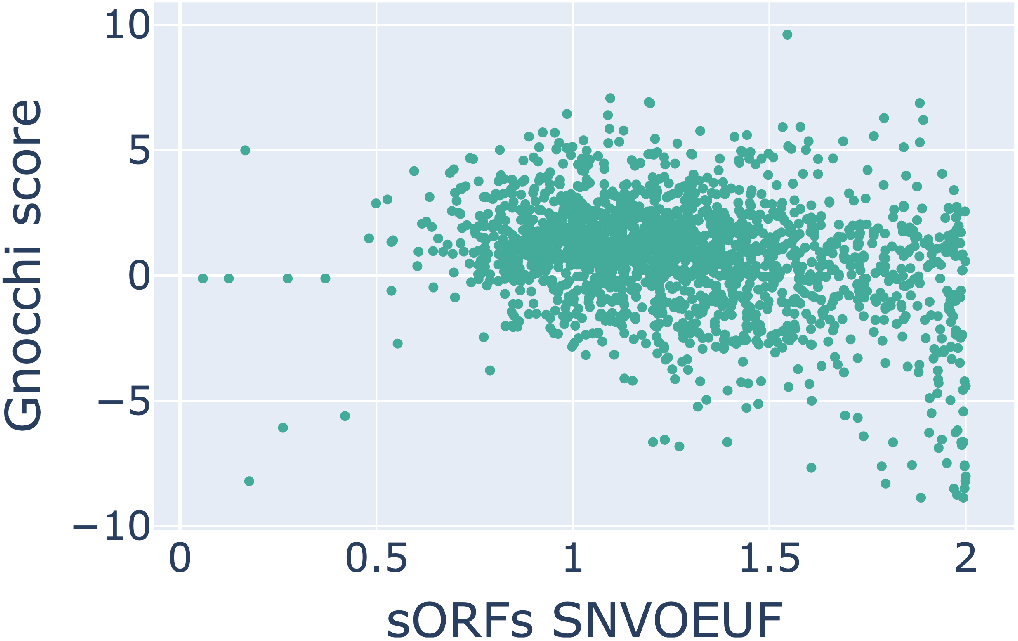
Comparison between SNVOEUF of sORFs located on lncRNAs calculated using gnomAD genomes and the Gnocchi Score.

**Fig. 29.**
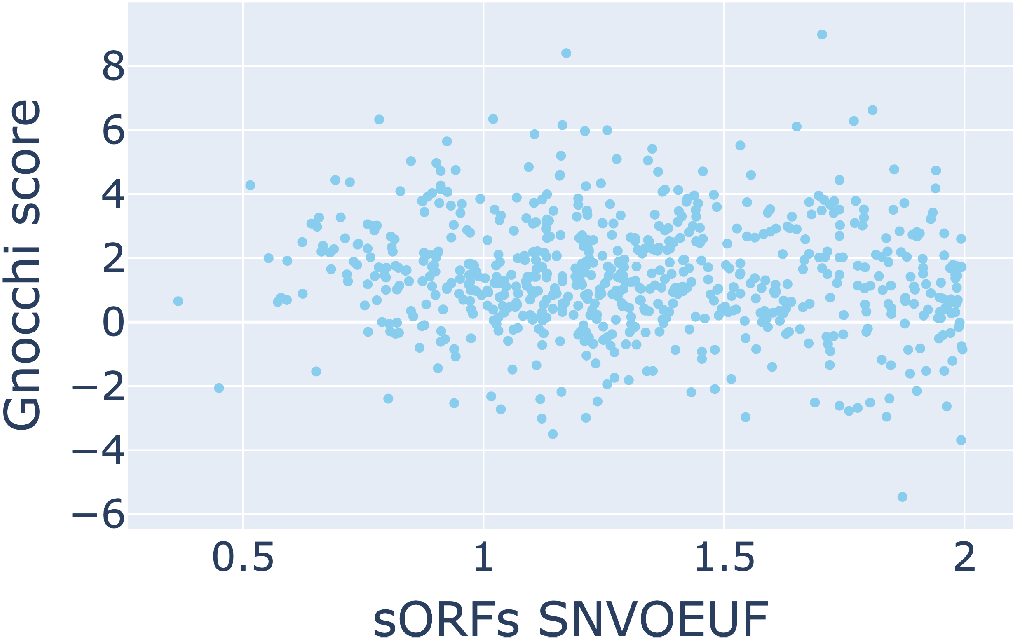
Comparison between SNVOEUF of uoORFs calculated using gnomAD genomes and the Gnocchi Score.

**Fig. 30.**
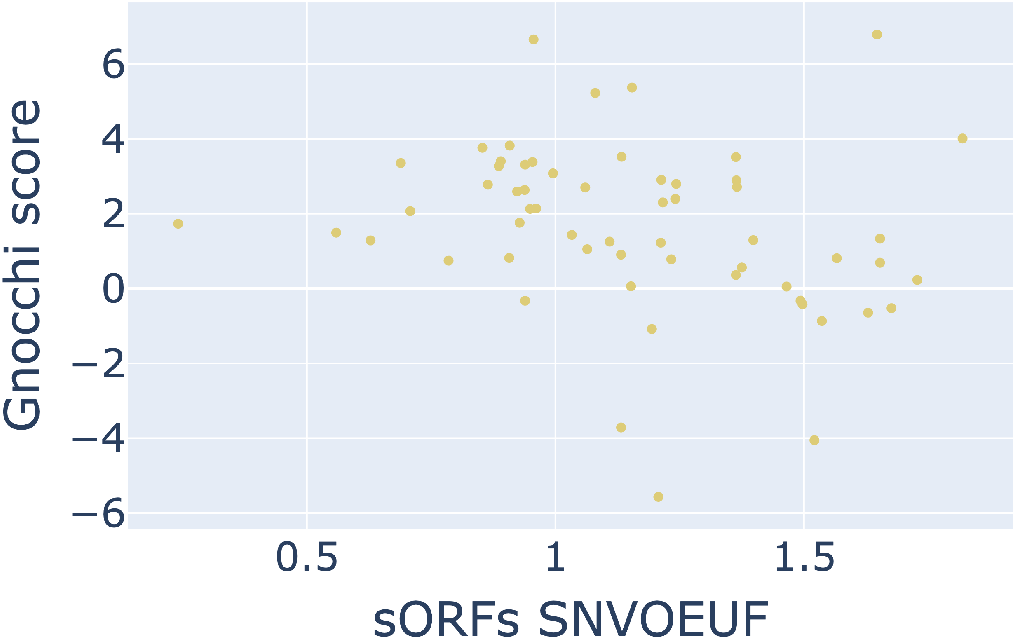
Comparison between SNVOEUF of doORFs calculated using gnomAD genomes and the Gnocchi Score.

**Fig. 31.**
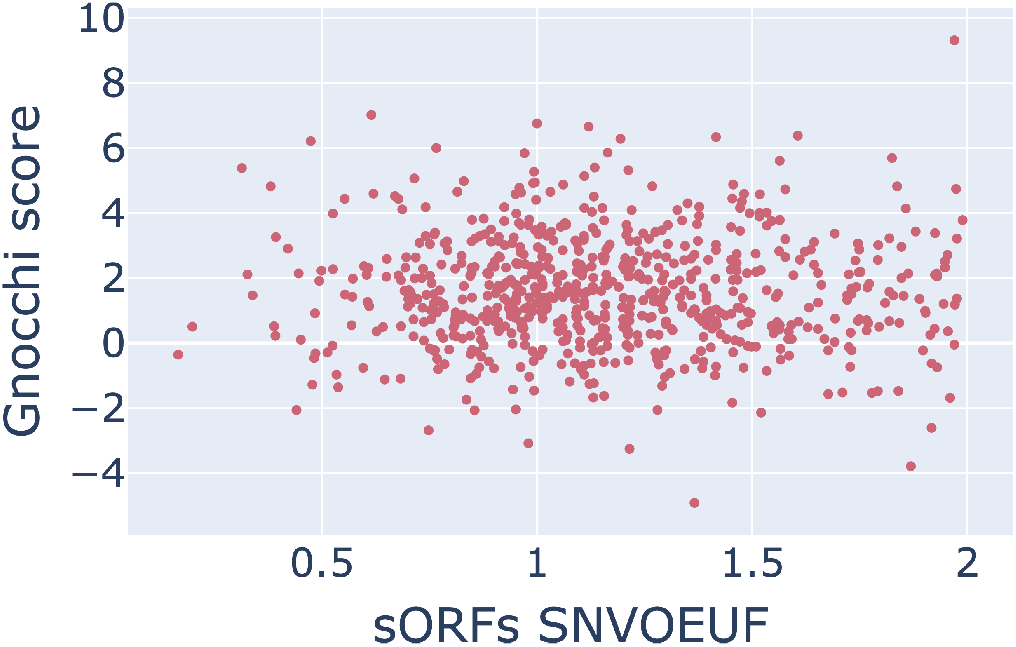
Comparison between SNVOEUF of intORFs calculated using gnomAD genomes and the Gnocchi Score.

### E. Individual sORF class comparison with the MOEUF of neighboured genes

Additionally, to the summary plot found in the article, we provide individual plots for comparison, in which we compare the MOEUF score of the individual sORF classes with the MOEUF scores of neighboured genes.

**Fig. 32.**
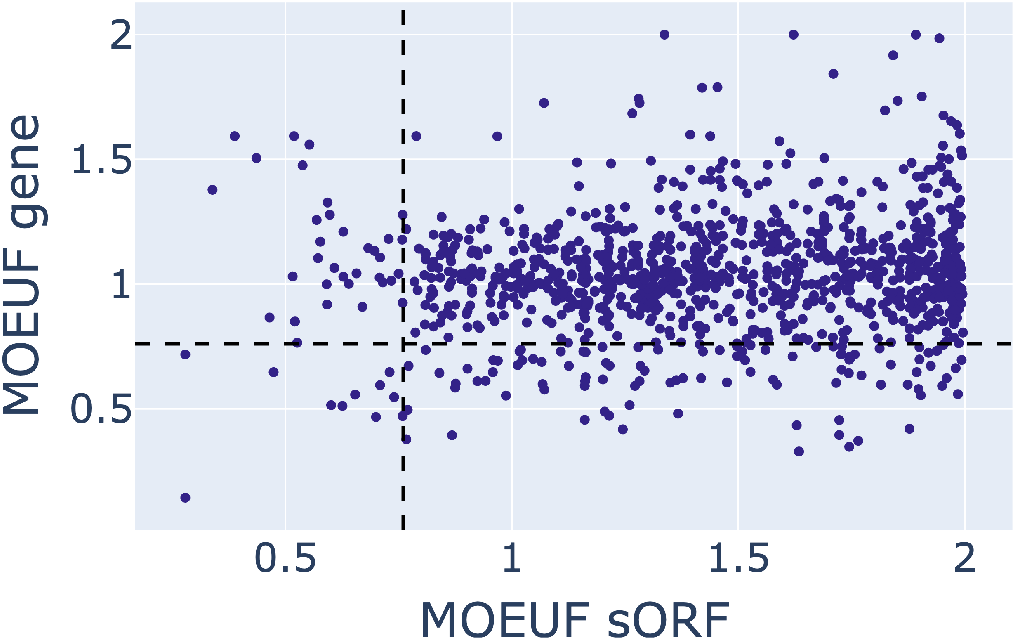
Comparison between MOEUF of uORFs and neighboured genes calculated using gnomAD genomes.

**Fig. 33.**
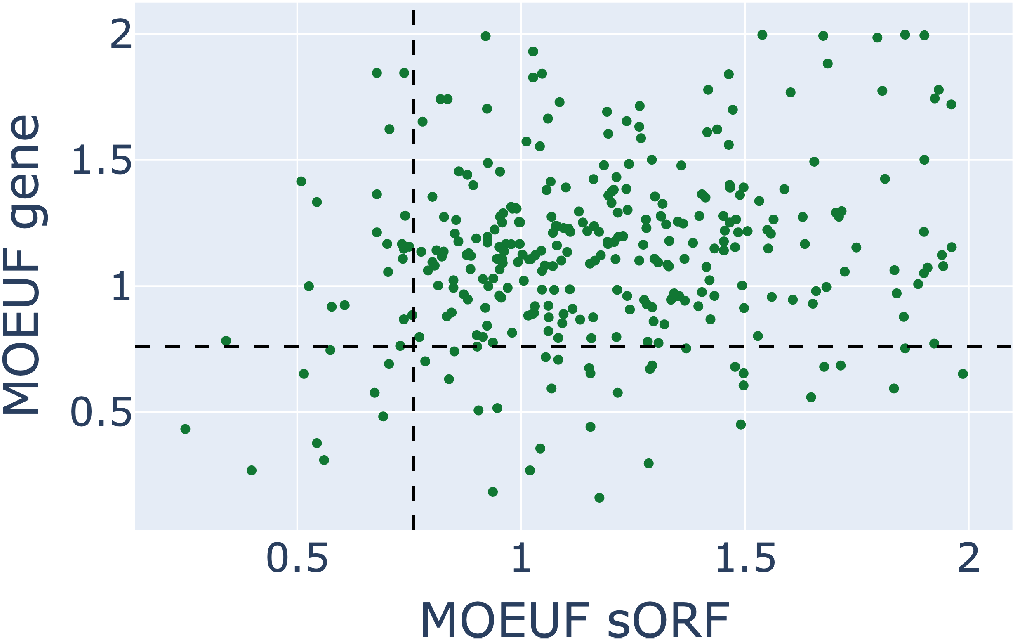
Comparison between MOEUF of dORFs and neighboured genes calculated using gnomAD genomes.

**Fig. 34.**
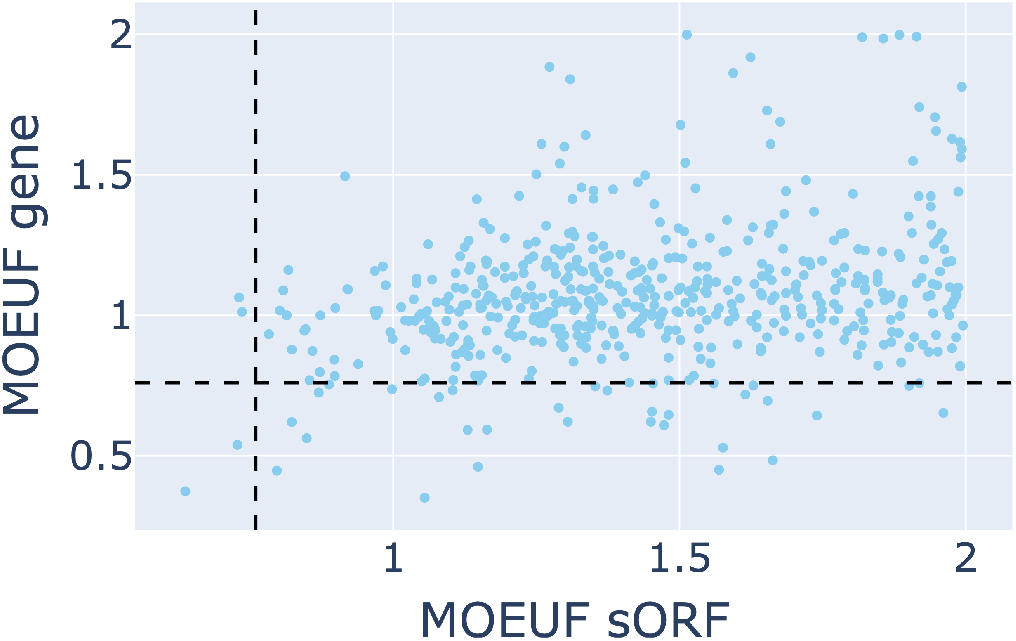
Comparison between MOEUF of uoORFs and neighboured genes calculated using gnomAD genomes.

**Fig. 35.**
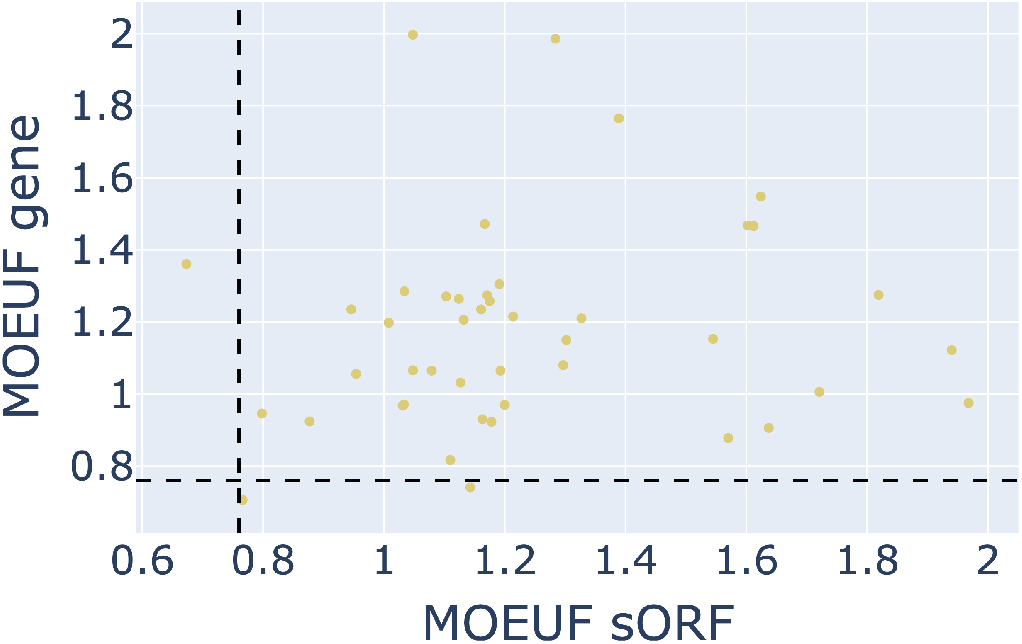
Comparison between MOEUF of doORFs and neighboured genes calculated using gnomAD genomes.

**Fig. 36.**
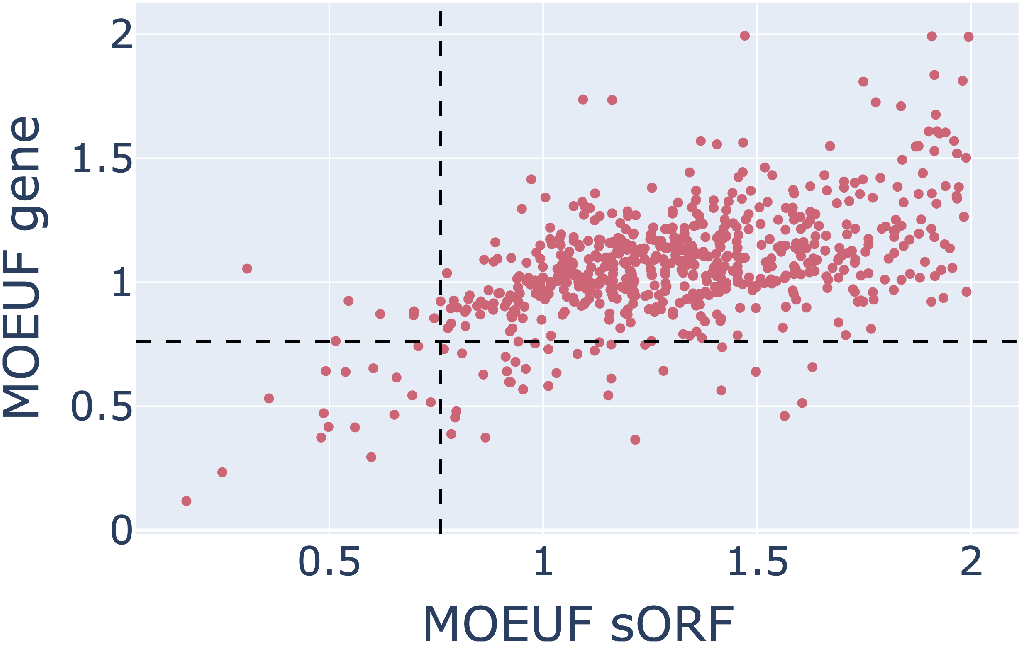
Comparison between MOEUF of intORFs and neighboured genes calculated using gnomAD genomes.

